# A non-tethering role for the Drosophila Pol θ linker domain in promoting damage resolution

**DOI:** 10.1101/2024.08.27.609911

**Authors:** Justin R. Blanch, Manan Krishnamurthy, Mitch McVey

## Abstract

DNA polymerase theta (Pol θ) is an error-prone translesion polymerase that becomes crucial for DNA double-strand break repair when cells are deficient in homologous recombination or non-homologous end joining. In some organisms, Pol θ also promotes tolerance of DNA interstrand crosslinks. Due to its importance in DNA damage tolerance, Pol θ is an emerging target for treatment of cancer and disease. Prior work has characterized the functions of the Pol θ helicase-like and polymerase domains, but the roles of the linker domain are largely unknown. Here, we show that the *Drosophila melanogaster* Pol θ linker domain promotes egg development and is required for tolerance of DNA double-strand breaks and interstrand crosslinks. While a linker domain with scrambled amino acid residues is sufficient for DNA repair, replacement of the linker with part of the *Homo sapiens* Pol θ linker or a disordered region from the FUS RNA-binding protein does not restore function. These results demonstrate that the linker domain is not simply a random tether between the helicase-like and polymerase domains. Furthermore, they suggest that intrinsic amino acid residue properties, rather than protein interaction motifs, are more critical for Pol θ linker functions in DNA repair.

## Introduction

DNA double-strand breaks (DSBs) are harmful DNA lesions that can be induced by exogenous agents such as ionizing radiation (IR) or normal cellular processes such as DNA replication and transcription (1). To prevent genomic catastrophe, metazoans frequently use two pathways to repair DSBs: non-homologous end joining (NHEJ) and homologous recombination (HR). NHEJ is favored when DNA resection is inhibited, involves limited processing and ligation of DSB ends, and can result in small mutations (2–4). HR requires DNA resection, strand invasion and fill-in synthesis, and is typically error-free (5). Since NHEJ and HR account for the majority of DSB repair, deficiencies in either pathway can lead to cell death or pathogenic mutations.

HR or NHEJ-deficient cells often utilize an alternative pathway for DSB repair and survival called theta-mediated end joining (TMEJ) (6–11). The most essential factor in the TMEJ pathway is DNA polymerase theta (Pol θ, *POLQ*). Like HR, TMEJ is promoted by DNA resection, but in contrast to HR, TMEJ regularly causes deletions and insertions in the genome (12–14). TMEJ-induced mutations and elevated levels of Pol θ frequently occur in cancers and have been associated with poor prognoses for cancer patients (13,15–21).

Pol θ possesses three domains: helicase-like, linker, and polymerase. The helicase-like and polymerase domains are largely conserved in metazoans, while linker domains are highly divergent (Figure 1A, Supplementary Figures S1 and S2). Prior work has shown that the helicase-like domain dissociates RAD51 and RPA from single-stranded DNA (ssDNA) and promotes microhomology synapsis needed for TMEJ (22–24). Following synapsis, the polymerase domain stabilizes synapsed ends and synthesizes new DNA to fill in gaps (25,26).

**Figure 1.**
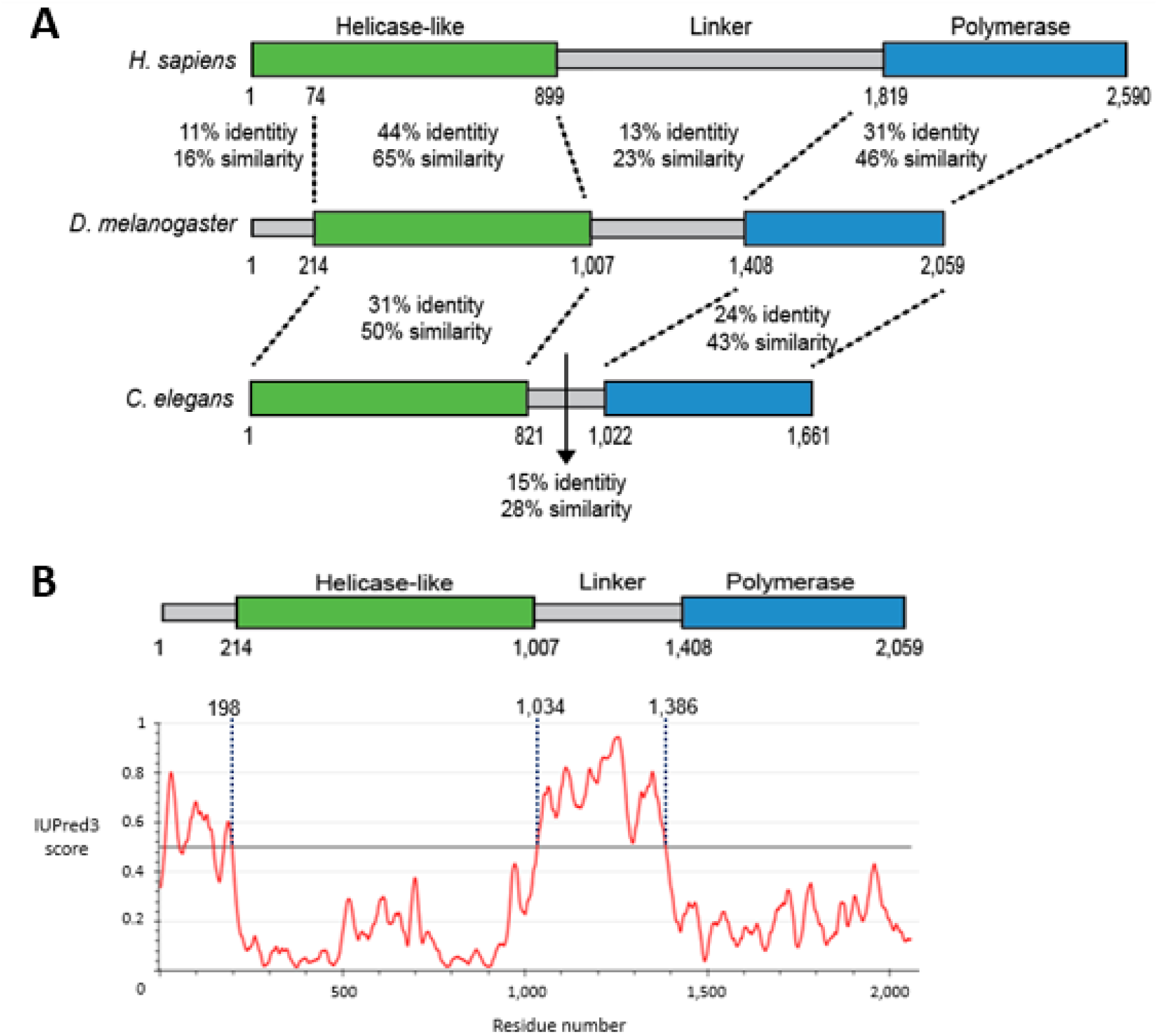
The Pol θ linker domain is mostly not conserved among metazoans and has a disordered structure. (**A**) Percent identity and similarity scores for pairwise alignments between Pol θ domains (Needleman-Wunsch global alignments, Blosum62 cost matrix, penalties: gap open=12, gap extension=3, similarity threshold=1). Alignments with previously defined *H. sapiens* helicase-like (1-899) and polymerase (1,819-2,590) domains (37,38) were used to determine domain boundaries in *D. melanogaster* and *C. elegans* sequences: domain ends coincide with the end of grouped identical residues (not shown, Geneious global alignments with free end gaps, Blosum62 cost matrix, penalties: gap open=12, gap extension=3). (**B**) Prediction of *D. melanogaster* Pol θ disorder using IUPred3 long disorder and strong smoothing tools (39). Residues with scores above the 0.5 threshold (horizontal line) are predicted to be disordered (40).

Mammalian cells deficient in Pol θ demonstrate sensitivity to IR-induced DNA breaks (27,28). HR-deficient Drosophila require Pol θ polymerase activity, but not helicase-like ATPase activity to resolve IR-induced breaks (29). In addition to repairing DSBs, Pol θ promotes resolution of interstrand crosslinks (ICLs) in *Mus musculus* embryonic fibroblasts, *Caenorhabditis elegans*, *Arabidopsis thaliana*, and *Drosophila melanogaster* (30–35). In Drosophila, both polymerase and helicase-like ATPase activities are required for organismal resistance to interstrand crosslinks (29). The mechanism for how Pol θ supports ICL tolerance is unknown, but *in vitro* evidence suggests Pol θ facilitates bypass of ICLs (29), possibly to encourage processive synthesis during DNA replication.

Roles of the Pol θ linker domain in DNA repair are largely unexplored. Previous reports showed that the *Homo sapiens* Pol θ linker domain is not required for TMEJ *in vitro* (22), but phosphorylated serine residues within the linker promote a Pol θ interaction with TOPBP1 and mitotic DSB repair *in vivo* (36). It is currently unclear how well the respective phosphorylation sites are conserved within Drosophila and other eukaryotes and if other regions in Pol θ linker domains may also influence DNA repair *in vivo*.

Pol θ linker domains contain intrinsically disordered regions (IDRs) (Figure 1B and Supplementary Figure S3). IDRs can encourage nuclear localization, cell cycle signaling, autoinhibition of protein activity, and protein-protein interactions (41,42). These functions are fine-tuned by post-translational modifications (PTMs) that alter domain architecture and respective enzyme activity or protein binding interfaces (43,44). Previous reports suggest IDRs influence the functions or recruitment of RAD52, p53, FUS, and several other DNA repair proteins (45–48). Accordingly, we postulated that the IDRs in the Pol θ linker domain may also be important for DNA repair.

To test this hypothesis, we performed mutagen sensitivity assays, measured egg hatching, and explored linker sequence requirements for repair using replacement IDRs. We found that the Pol θ linker domain is critical for repair of DNA double-strand breaks and tolerance to interstrand crosslinks and promotes normal Drosophila egg development. Interestingly, a randomly scrambled Drosophila linker, but not the *Homo sapiens* Pol θ linker sequence or a disordered region from the FUS protein, is sufficient for DNA damage tolerance. Overall, our findings establish that the linker domain is required for Pol θ-dependent repair of DNA damage *in vivo* and its functions are likely independent of amino acid residue arrangement.

## Materials and Methods

### Non-transgenic flies

All flies were raised at 25°C on a standard cornmeal diet with a 12:12 hour light/dark cycle. Wild-type flies were *w^1118^* (BDSC 3605) unless otherwise noted.

The *polq^3XFLAG,PBac{3XP3-DsRed}^* null allele contains a *piggyBac* transposon inserted by CRISPR-mediated knock-in at the endogenous *POLQ* locus (technique described in https://flycrispr.org/scarless-gene-editing/) (49,50). An 825 bp homology arm containing most of the *POLQ* promoter and an 800 bp homology arm containing most of *POLQ* exon 1 were amplified by PCR and inserted into pHD-w+ (Addgene 80927, gift from Kate O’Connor-Giles) (NEBuilder HiFi Assembly). BbsI restriction overhangs were incorporated on the ends of an oligo template (5’ GAAACAGCACCTTAATGGCGC 3’) for a gRNA that targets the beginning of the *POLQ* coding sequence. The gRNA oligo was inserted into pU6-Bbs1_chiRNA (Addgene 45946, gift from Melissa Harrison, Kate O’Connor-Giles, and Jill Wildonger) (51). The respective gRNA PAM site was mutagenized in a *POLQ* homology arm within pHD-w+ by Q5 site-directed mutagenesis (New England BioLabs). Another fragment containing *PBac{3XP3-DsRed}* was amplified (Addgene 80820, gift from Kate O’Connor-Giles) and inserted in between *POLQ* homology arms in pHD-w+. pHD-w+ and pU6-Bbs1_chiRNA were injected into *Cas9*-expressing embryos (BDSC 51324) to generate a *DsRed* transformant (BestGene). The *piggyBac* insertion created an early stop codon at the beginning of the endogenous *POLQ* locus and the *polq^3XFLAG,PBac{3XP3-DsRed}^* allele was validated by PCR (Primers 1 and 2 in Supplementary Table S1).

The *spn-A^5^* null mutant was made using CRISPR-induced mutagenesis. *Cas9* (BDSC 79006) and a *spn-A* gRNA (BDSC 76440) were expressed in the male germline. In a male offspring, a four-nucleotide deletion and early stop codon were recovered at the beginning of the *spn-A* coding sequence. The *spn-A^5^* allele was characterized by sequencing and validated in flies by PCR (Primers 3 and 4).

### Transgenic flies

To make the *polq^ΔL-untagged^* transgene, a 4.572 kb fragment, which included 1.287 kb upstream of the *POLQ* translation start site (endogenous *POLQ* promoter and 5’ UTR) and 3.252 kb downstream of the translation start site, was amplified using genomic DNA and primers containing BglII and NotI restriction sites. The restriction fragment was ligated into pattB (DGRC 1420). A 2.756 kb fragment, which included 2.329 kb upstream of the stop codon, 3’ UTR, and 242 bp downstream of the 3’ UTR, was ligated into pattB using primers with NotI and Acc65I restriction sites. 953 bp within exon 4 (the linker domain) were excluded from 5’ and 3’ *POLQ* restriction fragments. pattB-polq^ΔL-untagged^ was validated by sequencing and injected into *ΦC31*-expressing embryos (BestGene). Transgenes were inserted at *ZH-68E-attP* on chromosome 3L (BDSC 24485). *polq^ΔL-untagged^* was validated in Drosophila by PCR (Primers 6 and 7).

To make the 3xFLAG-tagged *polq^ΔL^* transgene, pattB-polq^ΔL-untagged^ and primers containing AvrII and SacI restriction sites were used to amplify 1.287 kb upstream of the *POLQ* translation start site. The restriction fragment was ligated into pAFW-Ago2 (Addgene 50554, gift from Yukihide Tomari) (52). Using pAFW-Ago2, a 1.395 kb fragment (promoter, 5’ UTR, Kozak sequence, and 3xFLAG) was amplified using primers with BamHI and BglII restriction sites and ligated into pattB. The *polq^ΔL-untagged^* transgene was amplified downstream of the translation start site using pattB-polq^ΔL-untagged^ as a template and primers with BglII and Acc65I restriction sites, then ligated into pattB. pattB-polq^ΔL^ was validated by sequencing and *polq^ΔL^* was validated in Drosophila by PCR (Primers 6 and 7).

pattB-POLQ^+^ was made in the same way as pattB-polq^ΔL^ except genomic DNA was used to amplify *POLQ*. pattB-POLQ^+^ was validated by sequencing and *POLQ^+^* was validated in Drosophila by PCR (Primers 5 and 2).

To make replacement-linker transgenes, gBlocks (Integrated DNA Technologies) were inserted into pattB-polq^ΔL^. The gBlocks coded for randomly scrambled *Drosophila melanogaster* linker residues 1,025-1,342, residues 1,483-1,800 from *Homo sapiens* Pol θ, and residues 2-214 from *Homo sapiens* FUS (Supplementary Table S2). NotI restriction sites and adapters for sequence stabilization were added to gBlock ends. Sequences were codon optimized for *Drosophila melanogaster* (Integrated DNA Technologies). gBlocks were ligated into pminiT2.0 for propagation (New England BioLabs PCR Cloning) and digested by NotI for insertion into pattB-polq^ΔL^ to make pattB-polq^SCR^, pattB-polq^HUM^, and pattB-polq^FUS^. gBlock orientation and composition within each plasmid were validated by PCR and sequencing (Primers 6 and 7). Transgenes were validated in Drosophila by PCR (*polq^SCR^*: primers 8 and 7, *polq^HUM^*: 6 and 9, *polq^FUS^*: 10 and 7).

### Quantitative PCR

To extract RNA, 9-15 second/third instar larvae or flies were homogenized in TRIzol (Thermo Fisher Scientific). The homogenate was vigorously mixed with chloroform and centrifuged (15 minutes, 12,000 x g at 4°C). RNA in the upper aqueous layer was removed, mixed with isopropanol, and incubated for 10 minutes at −20°C. After centrifugation (10 minutes, 12,000 x g at 4°C), the pellet was washed with 75% ethanol and allowed to dry. Pellets were resuspended in nuclease-free water and stored at −80°C.

A portion of each RNA sample was treated with DNase I in the respective buffer for 30 minutes at 37°. DNase I activity was inactivated in 4mM EDTA and by heating for 10 minutes at 72°C. DNaseI-treated 1 µg RNA aliquots were immediately used in cDNA synthesis and no-reverse transcriptase control (no-RT) reactions with random primers (New England BioLabs Protoscript II kit).

For qPCR reactions, cDNA and no-RT samples were diluted 1:3. The following reagents were loaded into each well: 10 µL of 2X SYBR Green Fast master mix (ABclonal), 0.4 µL of forward and reverse primers (0.2 µM of each), 7.2 µL of nuclease-free H_2_O, and 2 µL of the diluted sample. Biological replicates were run in triplicate for each genotype and primer set. One primer set amplified *POLQ*, but did not amplify *polq^3XFLAG,PBac{3XP3-DsRed}^* (Primers 11 and 12, Supplementary Figure S4 and Supplementary Table S3). The second primer set amplified an endogenous control gene, *rp49* (Primers 13 and 14). No-template controls for both primer sets were run in triplicate. Samples were analyzed in a QuantStudio 6 Flex Real-Time PCR system (Applied Biosystems) using the following cycling conditions: 50°C for 2 minutes, 95°C for 6 minutes, (95°C for 15 seconds, 60°C for 20 seconds) x 40 cycles, and a continuous melt curve stage (95°C for 15 seconds, 60°C for 20 seconds, 95°C for 15 seconds). *rp49* Ct triplicate averages were subtracted from *POLQ* Ct triplicate averages to calculate ΔCt for each biological replicate and calibrator ΔCt values were subtracted from biological replicate ΔCt values to calculate ΔΔCt and relative quantification (2^-ΔΔCt^).

RQmin and RQmax confidence intervals were calculated using the following adapted equations (Applied Biosystems):

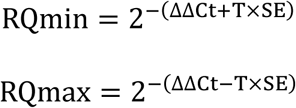

T=student’s two-tailed T factor for ΔCt mean where α=0.05 and DF=4.

SE=standard error of 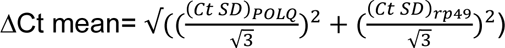 where Ct SD=triplicate standard deviation for a target.

### Immunoprecipitation and western blotting

*POLQ^+^*, *polq^null^*- or *polq^ΔL^*, *polq^null^*-homozygous flies were loaded into separate sets of 1.3-liter fly cages containing grape juice agar plates. 500 mg of embryos were collected for each genotype and frozen in liquid nitrogen. Embryos were suspended in lysis buffer (20 mM HEPES pH 7.6, 500 mM NaCl, 1.5 mM MgCl_2_, 10% glycerol, 0.1% Triton X-100, 1 mM phenylmethylsulfonyl fluoride, EDTA-free Pierce protease inhibitor (Thermo Fisher Scientific)) and sonicated for three 10 second bursts at 20% duty cycle with 10 second rests between bursts. Lysates were centrifuged to remove debris (4°C, 24,000 x g). For embryos carrying wild-type *POLQ* (BDSC 463), proteins were previously extracted using other established methods (53). Lysates were snap-frozen in liquid nitrogen and stored at −80°C.

For 3xFLAG immunoprecipitation, aliquots of ANTI-FLAG M2 Magnetic Beads (20 µL of packed gel volume per aliquot, MilliporeSigma) were washed twice with TBS (50 mM Tris-HCl, 150 mM NaCl) using a magnetic separator. 1 mL lysates were cleared by centrifugation (4°C, 14,000 x g) and added to the bead aliquots. Samples were rotated overnight at 4°C and supernatant was discarded. Beads were washed three times with TBS. FLAG-tagged protein was eluted twice using 100 µL of FLAG peptide (150 ng/µL, gift from Stephen Fuchs) during 30-minute rotations at 4°C.

Proteins were electrophoresed in a 4-20% SurePAGE gel and Tris-MOPS-SDS running buffer (GenScript). Proteins were transferred to a nitrocellulose membrane in wet conditions (25 mM Tris base, 25 mM bicine, 10% ethanol). The membrane was blocked (PBS, 5% non-fat milk) and probed with rabbit anti-DYDDDDK primary antibody (GenScript, 1:2,500 ; PBS-0.05% Tween 20, 1% non-fat milk) and rabbit IgG (HRP) secondary antibody (Abcam,1:10,000 ; PBS-0.05% Tween 20, 1% non-fat milk) with washes after antibody incubations (PBS-0.05% Tween 20). Horseradish peroxidase signal was induced using Prometheus ProSignal ECL (Genesee Scientific).

### Mutagen sensitivity assays

In IR assays, each cross included 20-60 females from one stock and 10-30 males from a separately maintained stock that were heterozygous for the same alleles. Crosses were housed at 25°C in 100 mL cages that included grape juice agar plates coated with yeast paste. Grape juice agar plates were replaced in cages every 24 hours. Once third instar larvae emerged, plates were either treated with Ce-137 radiation in a Gammator 1000 irradiator or left untreated. Larvae were transferred to bottles containing standard cornmeal food. Eclosed flies in each bottle were scored as homozygous or heterozygous and relative survival was calculated: ((homozygotes/total flies)_irradiated_ / (homozygotes/total flies)_untreated_ x 100). Progeny from 3-4 cross replicates were counted for each genotype. Crosses typically yielded hundreds of flies and not fewer than 32.

We anticipated that unmapped mutations, either randomly inherited during stock construction or caused by DNA repair deficiencies, could result in variable or misleading survival results in sensitivity assays. Accordingly, when possible, female and male parents were taken from separately constructed stocks of the same genotype to increase the likelihood that random mutations would be complemented in trans-heterozygous progeny. Reciprocal crosses, which contained reversed male and female stocks, were used to detect survival variation caused by uncomplemented random mutations on sex chromosomes in male progeny. Survival data for each reciprocal cross was analyzed separately (Supplementary Figure S6) before reciprocal data was combined for each genotype and dose (Figures 3 and 5).

In nitrogen mustard assays, each vial contained 5 females and 3 males heterozygous for specified alleles and followed the same complementation principles as IR assays. On day three, all crosses were transferred to a second set of vials. On day four, the first set of vials was treated with 0.007% nitrogen mustard. On day six, after flies were removed, the second set of vials was treated with water on day seven. Between 10 and 18 days after starting crosses, eclosed flies were scored as homozygous or heterozygous. Relative survival was not calculated for vials containing fewer than 10 flies. Progeny from 11-35 cross replicates were counted for each genotype.

### Hatching assays

Crosses included 8-30 females and 3-15 males homozygous for specified alleles. Crosses followed the same complementation principles as sensitivity assays and were housed at 25°C in cages that contained grape juice agar plates coated with yeast paste. Females were allowed to lay eggs on each plate for 24 hours. Hatched and unhatched eggs were counted 48 hours after each plate was removed from a cage. Eggs were counted from 3-6 crosses for each genotype and each cross produced greater than 44 eggs.

### Software

Graphs and statistical analyses were made using GraphPad Prism 10.1.2. Sequence analyses and alignments were made using Geneious Prime 2023.2.1. Accession numbers for alignment sequences are listed in Supplementary Table S4. Protein disorder graphs were made using IUPred3 (https://iupred3.elte.hu/).

## Results

### The Drosophila Pol θ linker domain is needed for DNA double-strand break repair and tolerance to interstrand crosslinks

To determine if the Pol θ linker domain promotes Drosophila resistance to DNA damage, we constructed a wild-type transgene and a *polq^ΔL^-*mutant transgene in which most of the disordered linker domain is deleted (Figure 2A). We designed the *ΔL* mutation to avoid removing partially conserved sequences near the helicase-like and polymerase domains. Both transgenes included 1.195 kb of the endogenous *POLQ* promoter. The transgenes were separately integrated into the left arm of chromosome 3 to create *polq^ΔL^* and *POLQ^+^* fly stocks. Each transgene was recombined onto a chromosome containing a *polq^null^* allele. *polq^ΔL^* and *POLQ^+^* transgenes are sufficiently expressed in larvae and adult flies relative to endogenous *POLQ* (Figure 2B, Supplementary Figure S4, and Supplementary Table S3) and the respective proteins can be detected via immunoprecipitation from Drosophila embryos (Figure 2C and Supplementary Figure S5).

**Figure 2.**
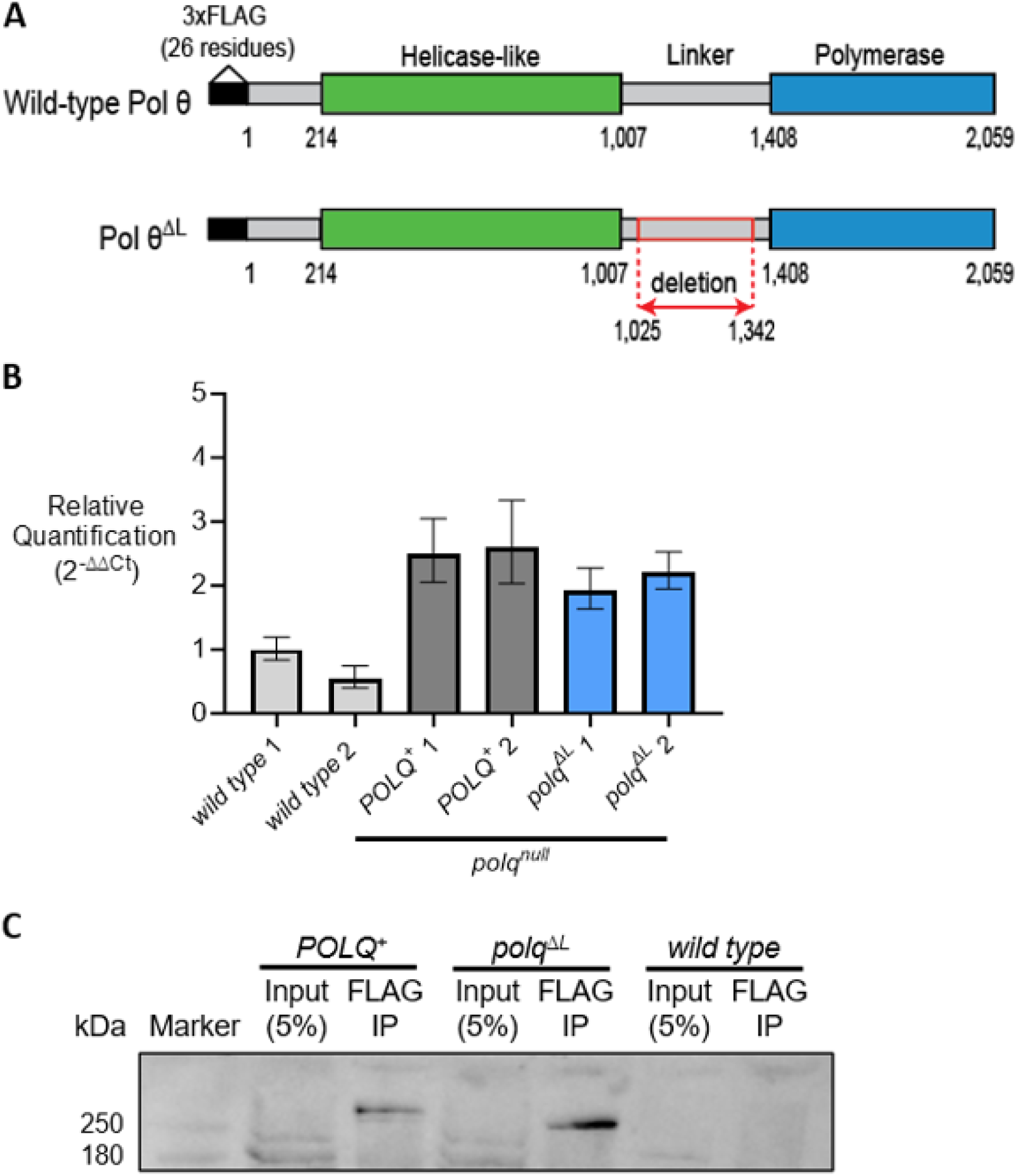
*POLQ* transgenes are expressed and their respective proteins are present *in vivo*. (**A**) Protein products from *POLQ^+^*and *polq^ΔL^* transgenes. Transgenes contain the endogenous *POLQ* promoter (not shown) and encode a N-terminal 3xFLAG tag. An in-frame deletion in *polq^ΔL^*subtracts linker residues 1,025-1,342. Transgenes were integrated on chromosome 3L and flies carried an endogenous *polq^null^* allele. (**B**) *POLQ* mRNA expression in larvae homozygous for endogenous *POLQ* (wild type) or *POLQ^+^, polq^null^* or *polq^ΔL^, polq^null^* alleles. 1 and 2 represent separately derived biological samples. Quantification was computed relative to endogenous control, *rp49* expression, and the calibrator sample, wild type 1. Error bars represent RQmin and RQmax confidence intervals (see methods). (**C**) Immunoprecipitated 3xFLAG-tagged Pol θ and Pol θ^ΔL^ from Drosophila embryo lysates. Proteins were detected using an anti-FLAG primary antibody.

We first analyzed roles of the linker domain in DSB repair. Previously, it was shown that HR-deficient Drosophila lacking the *Rad51* ortholog, *spn-A,* depend on Pol θ to repair IR-induced damage (54). Therefore, we put the Pol θ transgenic flies into a *spn-A* mutant background and measured IR sensitivity. We found that the wild-type *POLQ^+^* transgene provided similar tolerance to IR-induced DSBs as endogenous *POLQ* at doses of 50, 125, and 250 rads, but not 500 rads (Figure 3A, Supplementary Figure S6A, and Supplementary Table S5). Interestingly, *polq^ΔL^* mutants were hypersensitive to IR relative to *POLQ^+^* larvae and not significantly different from *polq^null^* mutants. These findings demonstrate that the linker domain is required for Pol θ-dependent DSB repair.

**Figure 3.**
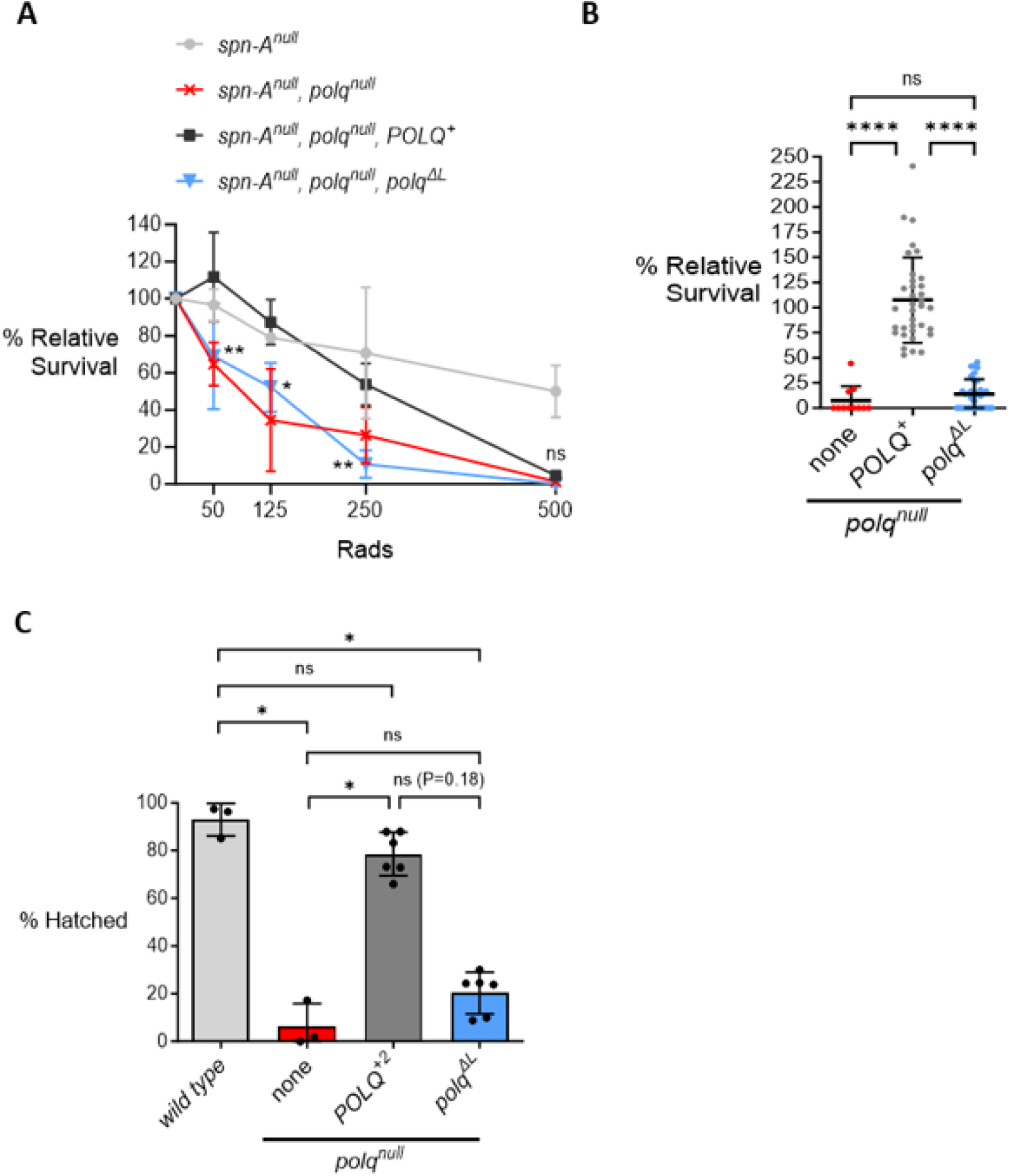
The linker domain is required for DNA damage tolerance and promotes egg hatching. (**A**) Survival of irradiated larvae to adulthood relative to untreated larvae. Each point represents mean relative survival from 3-4 replicate crosses and sample standard deviation is shown. Significance symbols denote comparisons between *spn-A^null^, polq^null^, polq^ΔL^* and *spn-A^null^, polq^null^, POLQ^+^*(two-way ANOVA and Sidak’s multiple comparisons test, all comparisons listed in Supplementary Table S5). (**B**) Mean relative survival of larvae treated with 0.007% nitrogen mustard from 11-35 crosses per genotype. Each point represents results for one cross. None=no transgene. Shown are sample standard deviation and comparisons using Kruskal-Wallis ANOVA and Dunn’s multiple comparisons test. (**C**) Mean egg hatching rates from 3-6 replicate crosses per homozygous genotype. Shown are standard deviations and statistical comparisons using Kruskal-Wallis ANOVA and Dunn’s multiple comparisons test. All panels: *p=0.01 to 0.05, **p=0.001 to 0.01, ****p<0.0001, ns=not significant.

In HR-competent Drosophila, Pol θ is critical for tolerance of ICLs induced by nitrogen mustard (29,30). We tested the ability of larvae lacking the Pol θ linker domain to survive to adulthood following nitrogen mustard treatment. We found that *polq^ΔL^* mutants were hypersensitive to nitrogen mustard and their survival was not significantly different from *polq^null^* (Figure 3B and Supplementary Figure S6B), indicating that the linker domain is required for either bypass or repair of ICLs.

We also wondered whether the linker domain is required for repair of endogenously derived DNA damage in Drosophila. Drosophila eggs lacking *POLQ* have thin eggshells and severe hatching defects, likely because they are unable to repair DNA breaks produced during gene amplification of the *DAFCs* (*Drosophila Amplicons in Follicle Cells*) (55–57). To assess if the Pol θ linker domain has a role in repair of these breaks, we measured hatching rates of eggs homozygous for *polq^ΔL^*. We compared *polq^ΔL^* eggs to *POLQ^+^* eggs from a new cross (*POLQ^+2^*), since *POLQ^+^* eggs had atypical hatching rates relative to other *POLQ^+^* stocks, possibly caused by randomly accumulated mutations (Supplementary Figures S6 and S7). *polq^ΔL^* eggs had substantially lower, albeit not significant, hatching frequencies than *POLQ^+2^* eggs and significantly lower hatching frequencies than wild-type eggs (Figure 3C and Supplementary Figure S6C). Interestingly, *polq^ΔL^* egg hatching rates were not significantly different from *polq^null^*, suggesting that the linker domain promotes resolution of endogenously derived DNA breaks during egg development.

We were curious if the N-terminal 3xFLAG tag in Pol θ^ΔL^ reduced the fitness of *polq^ΔL^* mutants in sensitivity and hatching assays. However, no significant differences were identified between *polq^ΔL^* and *polq^ΔL-untagged^* mutants (Supplementary Figure S7 and Supplementary Table S6). Taken together, our results strongly support that the linker domain is needed for Pol θ-dependent repair of DNA damage.

### Linker residue properties are important for Pol θ functions in DNA repair and damage tolerance

Since Pol θ linkers are not well-conserved among Drosophila species (Supplementary Figure S2), we hypothesized that amino acid residue properties may be more critical for linker functions than protein-binding motifs or catalytic subdomains. At the primary sequence level, Pol θ linker domains are disordered and composed mostly of hydrophilic polar or charged residues (Table 1, Supplementary Figures S3 and S8).

**Table 1.** Linker domain properties within Pol θ homologs.

| Organism | Predicted boundaries | Length (residues) | Overall charge | % Polar uncharged | % Charged | % Hydrophobic |
| --- | --- | --- | --- | --- | --- | --- |
| *H. sapiens* | 900-1,818 | 919 | -16 | 36 | 27 | 37 |
| *R. norvegicus* | 898-1,777 | 880 | -27 | 34 | 28 | 39 |
| *M. musculus* | 896-1,773 | 878 | -18 | 34 | 27 | 39 |
| *D. rerio* | 1,101-1,827 | 727 | -38 | 35 | 31 | 35 |
| *C. elegans* | 822-1,021 | 200 | -23 | 26 | 36 | 39 |
| *A. thaliana* | 1,320-1,601 | 282 | -7 | 31 | 25 | 46 |
| *D. melanogaster* | 1,008-1,407 | 400 | +2 | 35 | 26 | 39 |
| *D. melanogaster* helicase-like | 214-1,007 | 794 | +1 | 25 | 25 | 51 |
| *D. melanogaster* polymerase | 1,408-2,059 | 652 | +14 | 29 | 25 | 47 |

Pairwise alignments with human helicase-like and polymerase domains were used to predict linker boundaries. *D. melanogaster* Pol $\theta$ helicase-like and polymerase domains are listed for comparison to disordered linker domains. Properties were calculated in Geneious Prime. Charge was determined at pH 7.

To explore residue properties of the linker domain that are required for repair, we designed three replacement IDRs (Table 2 and Figure 4A). The first consists of randomly scrambled *D. melanogaster* linker residues. Length, disorder, overall charge, hydrophilic distribution, and residue composition are similar between the scrambled and wild-type linkers, while potential protein-binding motifs are likely interrupted in the scrambled sequence (Table 2 and Supplementary Figures S8, S9, and S10). The second includes part of the human Pol θ linker domain that is also disordered and has a similar length, hydrophilic distribution, and residue composition as the Drosophila linker (Table 2 and Supplementary Figures S8, S9, and S10). Notably, the human linker has an overall negative charge that contrasts with the positive charge of the Drosophila linker. The third consists of the N-terminal low-complexity domain from *Homo sapiens* FUS, which promotes aggregation at DNA damage for NHEJ and HR repair (58–60). The FUS low-complexity domain is highly disordered (Supplementary Figure S10), but has fewer residues and is more polar and hydrophilic than the Drosophila linker (Table 2, Supplementary Figures S8 and S9).

**Figure 4.**
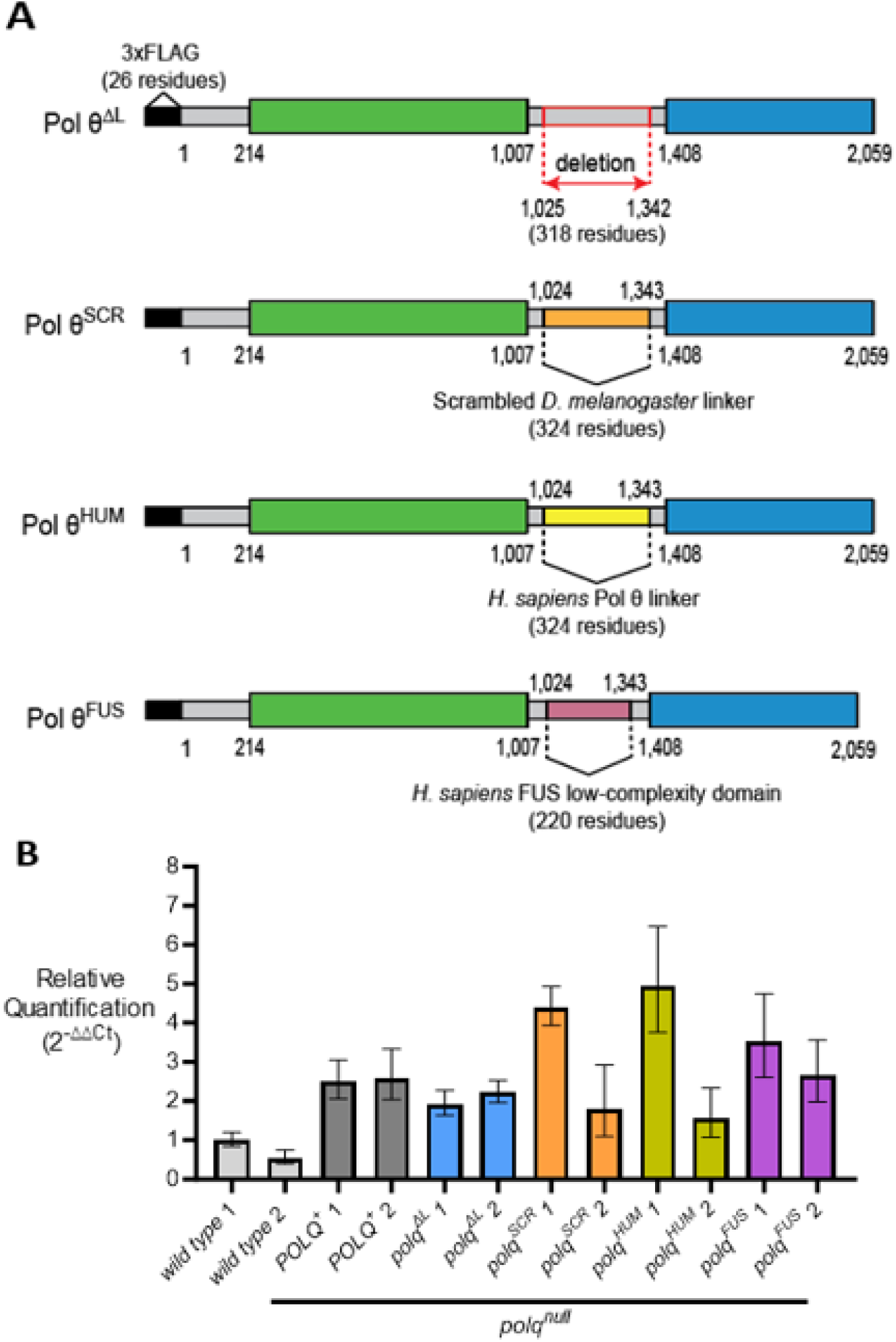
*polq*-mutant transgenes with replacement linkers are expressed *in vivo*. (**A**) The deleted linker in Pol θ^ΔL^ was replaced with the *D. melanogaster* linker sequence scrambled in a random order (Pol θ^SCR^), or residues 1483-1800 from the linker domain of human Pol θ (Pol θ^HUM^), or residues 2-214 from the N-terminal low-complexity domain of human FUS (Pol θ^FUS^) (58,59). Transgenes were integrated at the same chromosome *3* site as *polq^ΔL^* and put into a *polq^null^* background. (**B**) *POLQ* mRNA expression in wild-type larvae or those homozygous for a transgene and *polq^null^*. 1 and 2 represent separately derived biological samples. The endogenous control, *rp49* expression, and the calibrator sample, wild type 1, were used to calculate relative quantification. Error bars represent RQmin and RQmax. Wild-type, *POLQ^+^*, and *polq^ΔL^* data are also shown in Figure 2B.

**Table 2.** Replacement-linker properties.

| ***D. melanogaster***<br>**Pol θ linker** | **Length**<br>**(residues)** | **Overall**<br>**charge** | **% Polar**<br>**uncharged** | **%**<br>**Charged** | **%**<br>**Hydrophobic** |
| --- | --- | --- | --- | --- | --- |
| Wild type (deleted<br>in $\Delta L$ mutant) | 318 | +7 | 35 | 27 | 39 |
| Scrambled (SCR) | 324 | +7 | 34 | 26 | 40 |
| Human (HUM) | 324 | -26 | 33 | 27 | 40 |
| FUS (FUS) | 220 | -4 | 63 | 3 | 35 |

Replacement linker DNA sequences were codon optimized for Drosophila and separately inserted at the deletion site in *polq^ΔL^* to make new transgenes that were integrated into the same chromosomal location as the *polq^ΔL^* transgene. Replacement-linker transgenes had similar expression patterns as *polq^ΔL^* and *POLQ^+^* transgenes (Figure 4B, Supplementary Figure S4, and Supplementary Table S3).

In a *spn-A* null mutant background, we observed that *polq^SCR^* and *POLQ^+^* larvae had similar sensitivities to IR, while *polq^HUM^* and *polq^FUS^* mutants were hypersensitive (Figure 5A and Supplementary Table S7). Interestingly, *polq^SCR^* mutants were significantly more resistant than *POLQ^+^* larvae at 250 rads and *polq^HUM^* and *polq^FUS^* mutants were significantly more sensitive than *polq^ΔL^* mutants at 125 rads. These observations suggest that the scrambled linker promotes efficient Pol θ-dependent DSB repair, while human and FUS linkers are not sufficient and may even inhibit DSB repair.

**Figure 5.**
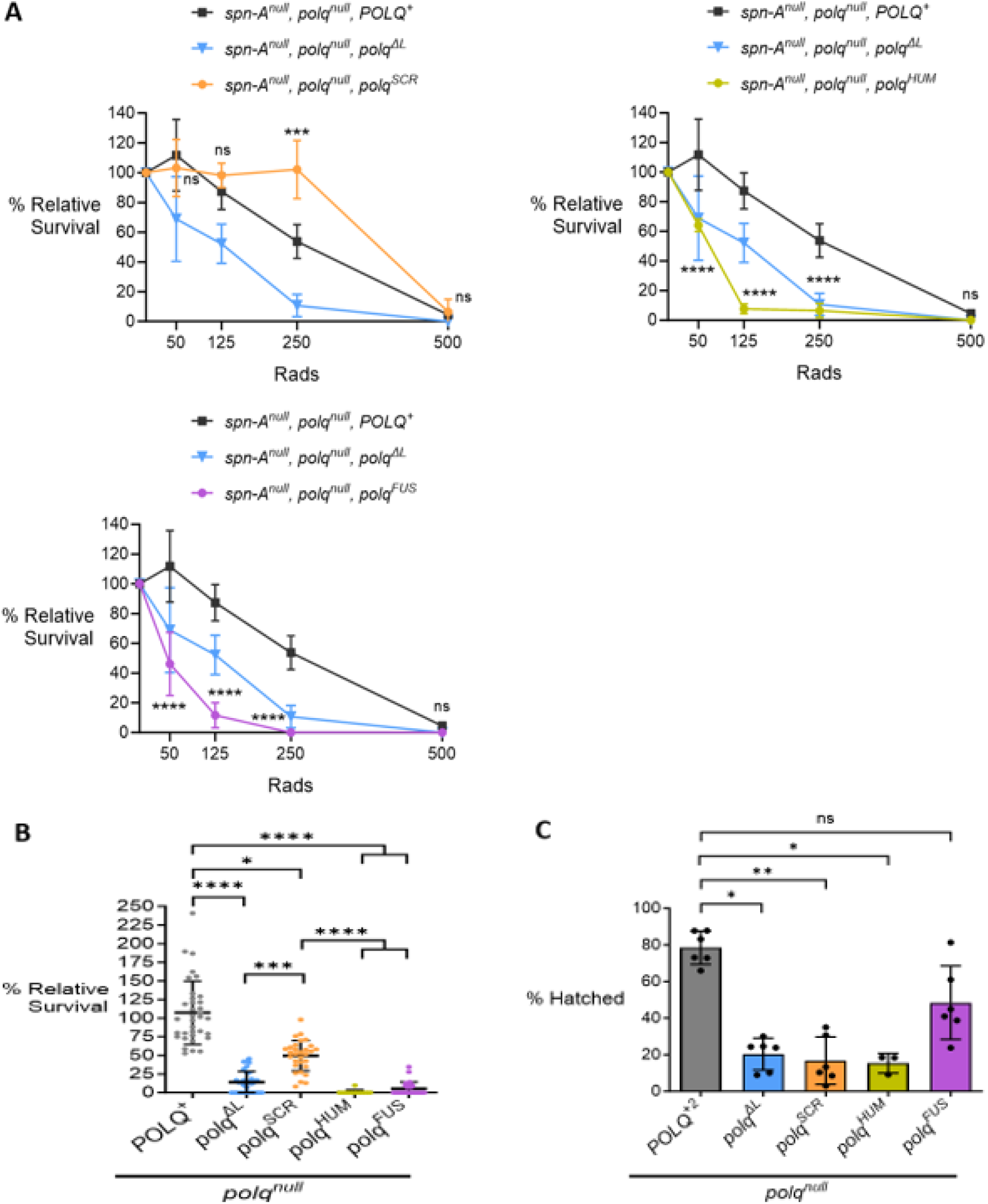
The scrambled linker rescues Pol θ-dependent resolution of DNA damage and the FUS linker rescues egg hatching. (**A**) Relative survival of irradiated larvae to adulthood. Each point represents mean survival from 3-4 crosses and sample standard deviation is shown. Data for *spn-A^null^, polq^null^, polq^ΔL^*and *spn-A^null^, polq^null^, POLQ^+^* are also shown in Figure 3A. Significance symbols denote comparisons between *spn-A^null^, polq^null^, POLQ^+^* and replacement-linker mutants (two-way ANOVA and Sidak’s multiple comparisons test, all comparisons shown in Supplementary Table S7). (**B**) Relative survival of larvae treated with 0.007% nitrogen mustard. Shown are mean survival from 12-35 crosses per genotype and sample standard deviation. Data for *polq^null^, POLQ^+^* and *polq^null^, polq^ΔL^* are also shown in Figure 3B. Nonsignificant comparisons are not shown (Kruskal-Wallis ANOVA and Dunn’s multiple comparisons test). (**C**) Mean hatching rates of eggs homozygous for transgenes and *polq^null^*. Standard deviation is shown for 3-6 crosses per genotype. Each cross consisted of two parental fly stocks homozygous for a particular genotype, except for *polq^null^, polq^HUM^* which was crossed with itself (see methods and Supplementary Figure S6). Data for *polq^null^, POLQ^+2^* and *polq^null^, polq^ΔL^* are also shown in Figure 3C. Most nonsignificant comparisons are not shown (Kruskal-Wallis ANOVA and Dunn’s multiple comparisons test). All panels: *p=0.01 to 0.05, **p=0.001 to 0.01, ***p=0.0001 to 0.001, ****p<0.0001, ns=not significant.

Following treatment with nitrogen mustard, the *polq^SCR^* transgene rescued survival relative to the *polq^ΔL^* transgene, although not to the level of the wild-type transgene (Figure 5B). *polq^HUM^* and *polq^FUS^* mutants rarely survived and were not significantly different from *polq^null^* mutants. This indicates that, in addition to DSB repair, the scrambled linker can promote tolerance to interstrand crosslinks, while the human and FUS linkers are unable to substitute for this function.

Lastly, we measured hatching rates of eggs homozygous for replacement-linker transgenes. Surprisingly, *polq^FUS^*-mutant hatching was not significantly different from *POLQ^+2^* hatching while *polq^SCR^* and *polq^HUM^* eggs had low hatching frequencies (Figure 5C). These findings suggest that the scrambled domain sequence is unable to support normal egg development, while the FUS low-complexity domain may share properties with the Drosophila linker that are required for DNA repair during egg development.

## Discussion

Pol θ-mediated DNA repair is critical for genome maintenance and cellular survival. Pol θ promotes tolerance of exogenously induced ICLs and is often required for resolution of DSBs in NHEJ- and HR-deficient cells. However, TMEJ frequently causes insertions and deletions in the genome that can elicit human disease. Cancer cells with a variety of genetic backgrounds require Pol θ to survive, but comprehensive roles of Pol θ domains in cell survival mechanisms are not entirely understood.

While prior investigations identified multiple functions for the Pol θ helicase-like and polymerase domains, the linker domain has been largely ignored. In this study, we have shown that the Drosophila Pol θ linker domain is required for DNA damage tolerance and has a specific sequence composition that promotes its functions. Importantly, flies possessing a deletion of the disordered linker region are unable to support any of the functions of wild-type Pol θ that we tested, including DSB repair, ICL tolerance, and egg viability.

### Functions of the linker domain in DSB and ICL repair

The importance of the linker domain in DSB repair was initially surprising, given it is disordered, mostly not conserved in Drosophila species, and the human linker domain is largely dispensable for *in vitro* (22) and *in vivo* DSB repair (61). Since Pol θ linkers are mostly not conserved, we postulated that the linker functions may depend on residue composition instead of binding motifs or catalytic subdomains. We explored this hypothesis by designing replacement linkers that have similar properties to the Drosophila linker domain. Interestingly, the scrambled *Drosophila melanogaster* linker fully rescued DSB repair and substantially rescued ICL tolerance. In contrast, part of the human Pol θ linker domain and the entire FUS low-complexity domain did not rescue survival in sensitivity assays. Since the human and FUS replacement linkers did not rescue repair, we think it is unlikely that the linker is simply acting as a nonspecific tether between the helicase-like and polymerase domains.

Damage tolerance promoted by the scrambled linker suggests that the ratio of linker residues is important, while linker-mediated protein interactions are unlikely needed for resolution of exogenously induced DNA damage in Drosophila. Prior work showed that the human Pol θ linker interacts with TOPBP1 to promote mitotic DSB repair (36). It has also been reported that the human Pol θ linker interacts with RAD51 to suppress HR (6), although this remains to be validated (35,62,63). At the moment, it is unclear if Drosophila Pol θ linker domains interact with other proteins. Protein interactions may be more common in mammalian Pol θ linker domains since they are more conserved than Drosophila linker domains (Supplementary Figure S1 and S2). However, some regions in Drosophila linker domains are partially conserved and may interact with proteins in specific DNA repair or developmental contexts to promote Pol θ functions.

Based on our findings, we propose that the Drosophila Pol θ linker could promote damage tolerance in at least two ways. First, the slightly positive, but overall neutral charge of the linker may allow interactions between the negatively charged N-terminus and positively charged polymerase domain (Supplementary Figure S11). These interactions could be critical for the helicase-like domain to be situated near the polymerase domain during microhomology synapsis and subsequent DNA synthesis. The scrambled linker may have rescued repair because it has the same overall charge and polarity as the Drosophila linker, while the strong negative charge of the human linker and strong polarity of the FUS linker could have disrupted Pol θ termini interactions.

Second, the linker domain may regulate Pol θ multimerization at DNA damage. When *in vitro* and not bound to DNA, the human linker suppresses Pol θ from forming multimers that promote TMEJ (22). The wild-type and scrambled linkers may function similarly *in vivo* and prevent unnecessary Pol θ multimerization prior to DNA damage.

### Potential roles of the Pol θ linker domain in Drosophila egg development

Re-replication of the *DAFC* genomic regions within Drosophila follicle cells is beneficial for eggshell development, but also causes DNA breaks, some of which require Pol θ for repair (55,56). Like *polq^null^* mutants in our study, *polq^ΔL^* eggs had low hatching frequencies. Surprisingly, *polq^SCR^* and *polq^HUM^* eggs also had low hatching frequencies while the *polq^FUS^* transgene partially rescued hatching. Since the *polq^FUS^* transgene did not rescue tolerance in sensitivity assays, repair of re-replication-induced DNA damage and exogenously induced damage likely require different linker-mediated mechanisms.

The FUS low-complexity domain normally encourages FUS aggregation at DNA damage (58–60), which is promoted by PARylation and discouraged by phosphorylation (64,65). Thus, one possibility is that FUS PTMs mimic Drosophila linker PTMs, which are required for Pol θ-mediated DNA repair during re-replication but are not sufficient to repair exogenously induced damage.

## Conclusion

Overall, we establish that residue specificity plays an important role in how the linker domain regulates Pol θ-dependent DNA damage tolerance in Drosophila. Our findings highlight that future studies should consider how the linker domain may influence overall Pol θ functions and the efficacy of Pol θ inhibitors for cancer treatment.

## Data Availability

Primary data that underlie the main and supplementary figures can be found in the supplementary Excel file.

## Supplementary Data

Supplementary Data are available at NAR Online.

## Acknowledgements

We thank Anna Joseph for creating the *spn-A^5^* mutant and Alice Miller for assistance with embryo lysis and fly counting.

## Author Contributions

J.R.B. and M.M. conceptualized and managed the study. J.R.B designed the methodology, performed the experiments, analyzed the data, and validated reproducibility of the results. M.K. designed and created the *polq^3XFLAG,PBac{3XP3-DsRed}^* allele. J.R.B. wrote the paper. J.R.B., M.M., and M.K. reviewed the paper and provided edits.

