## Supplemental figures and tables for "A non-tethering role for the Drosophila Pol θ linker domain in promoting damage resolution"

### A non-tethering role for the Drosophila DNA polymerase theta linker domain in promoting damage resolution

#### Supplementary Data

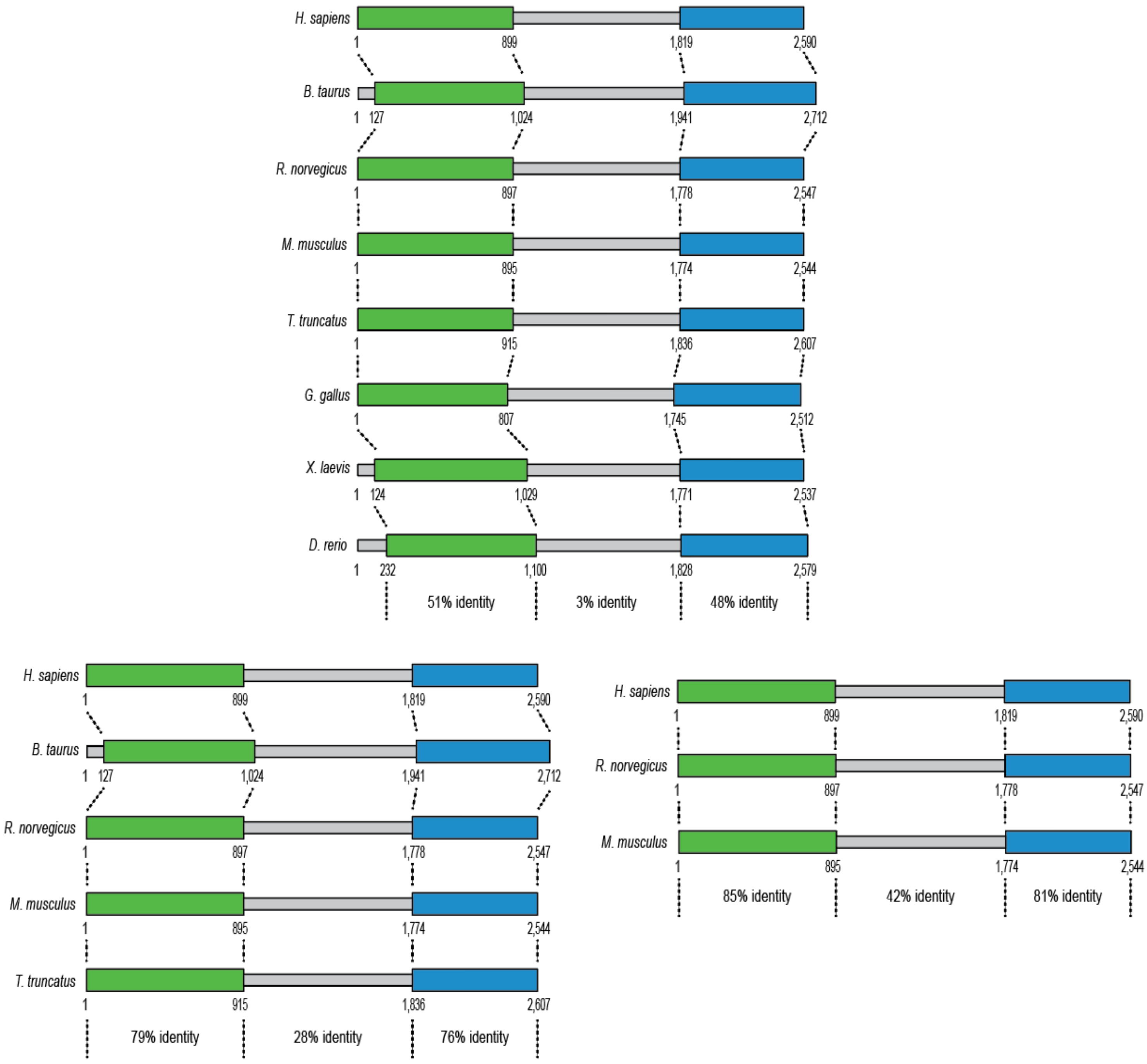


**Figure S1.** The Pol θ linker domain is mostly not conserved among vertebrates. Percent identity for multiple alignments between Pol θ domains (Geneious global alignments, no free end gaps, Blosum62 cost matrix; penalties: gap open=12, gap extension=3; refinement iterations=2). Previously mentioned alignments with human helicase-like and polymerase domains were used to predict protein domain boundaries. Each panel includes a separate alignment for each domain.

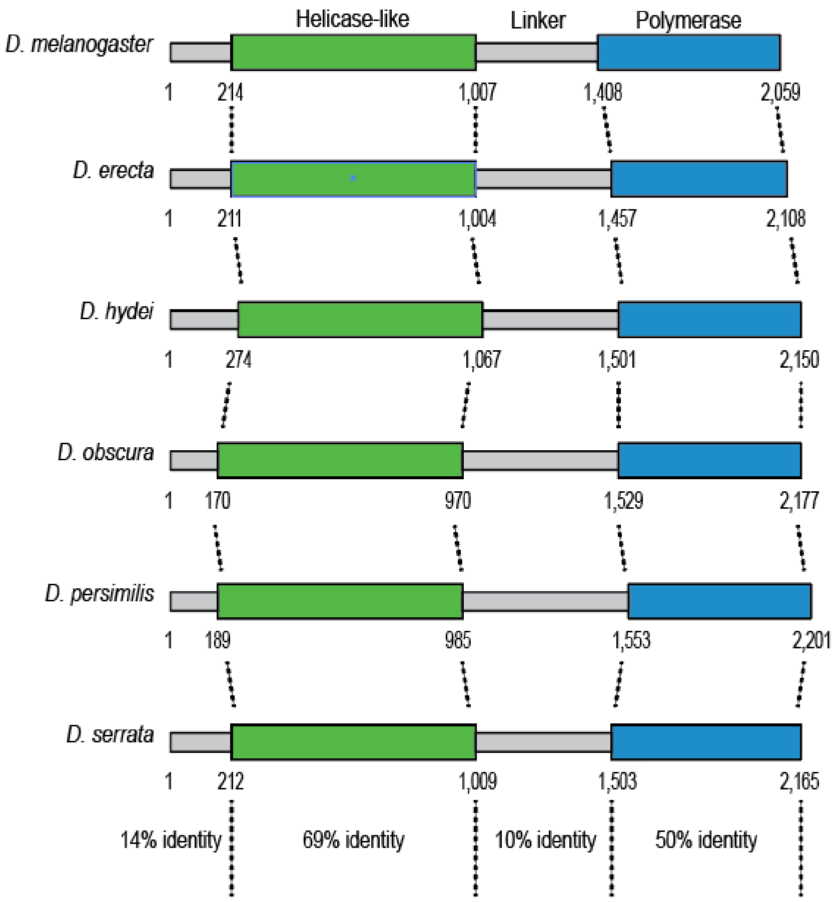


Pairwise comparisons with *D.melanogaster*

| | N-term | | Helicase-like | | Linker | | Polymerase | |
|---|---|---|---|---|---|---|---|---|
| | % identity | % similarity | % identity | % similarity | % identity | % similarity | % identity | % similarity |
| *D. erecta* | 69 | 79 | 89 | 95 | 57 | 67 | 83 | 93 |
| *D. hydei* | 32 | 46 | 80 | 90 | 30 | 44 | 71 | 85 |
| *D. obscura* | 32 | 47 | 81 | 90 | 23 | 35 | 64 | 79 |
| *D. persimilis* | 32 | 48 | 79 | 90 | 22 | 34 | 66 | 81 |
| *D. serrata* | 51 | 63 | 85 | 93 | 35 | 50 | 74 | 87 |

**Figure S2.** The linker domain is mostly not conserved among Drosophila species. Percent identity for domain multiple alignments among 6 Drosophila Pol θ homologs (Geneious global alignments, no free end gaps, Blosum62 cost matrix; penalties: gap open=12, gap extension=3; refinement iterations=2). Tables contain identity and similarity scores for pairwise alignments between *Drosophila melanogaster* Pol θ and homolog domains (similarity threshold=1).

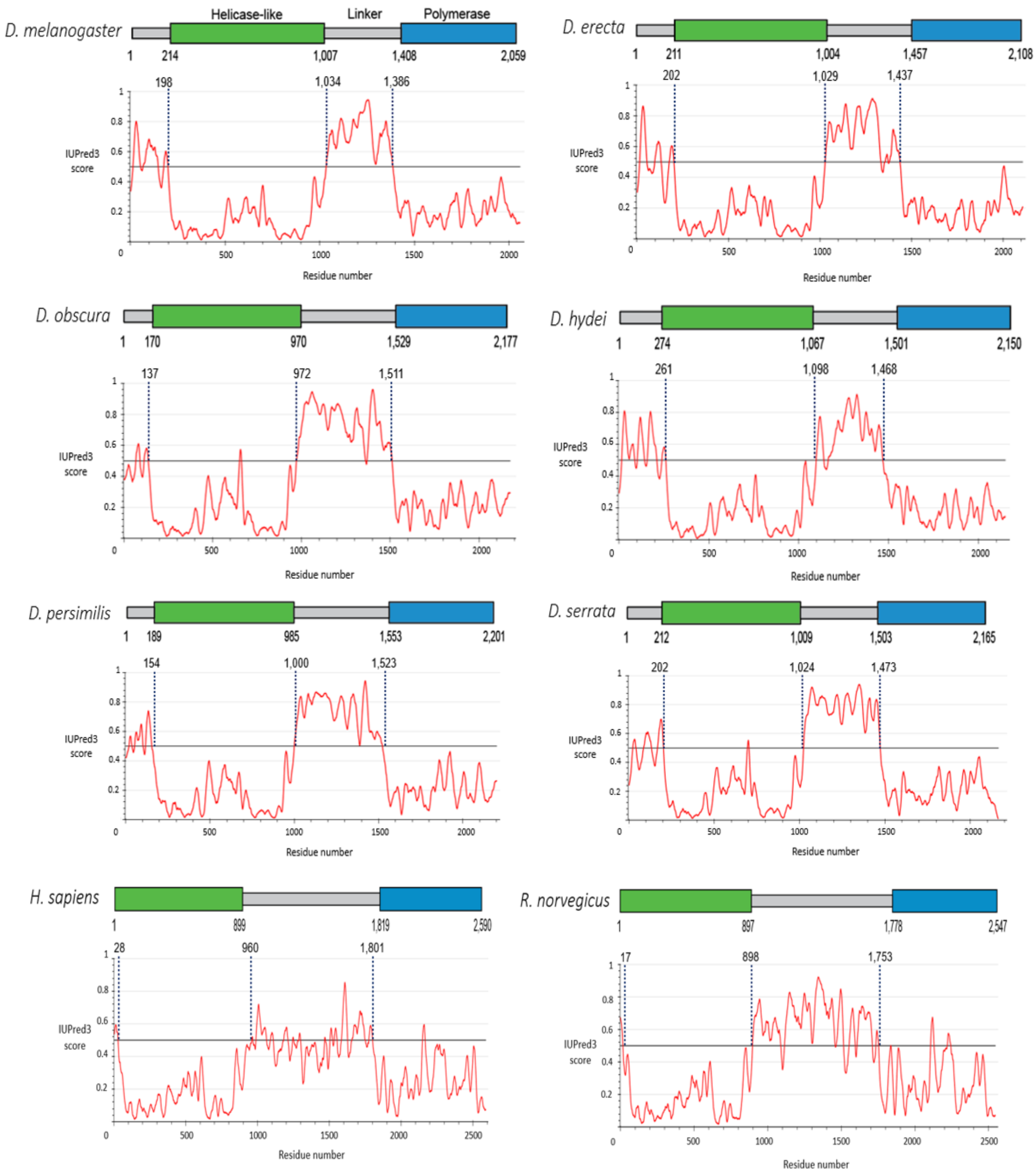

D. melanogaster
Helicase-like
Linker
Polymerase
1
214
1,007
1,408
2,059
198
1,034
1,386
IUPred3 score
Residue number
D. erecta
1
211
1,004
1,457
2,108
202
1,029
1,437
D. obscura
1
170
970
1,529
2,177
137
972
1,511
D. hydei
1
274
1,067
1,501
2,150
261
1,098
1,468
D. persimilis
1
189
985
1,553
2,201
154
1,000
1,523
D. serrata
1
212
1,009
1,503
2,165
202
1,024
1,473
H. sapiens
1
899
1,819
2,590
28
960
1,801
R. norvegicus
1
897
1,778
2,547
17
898
1,753

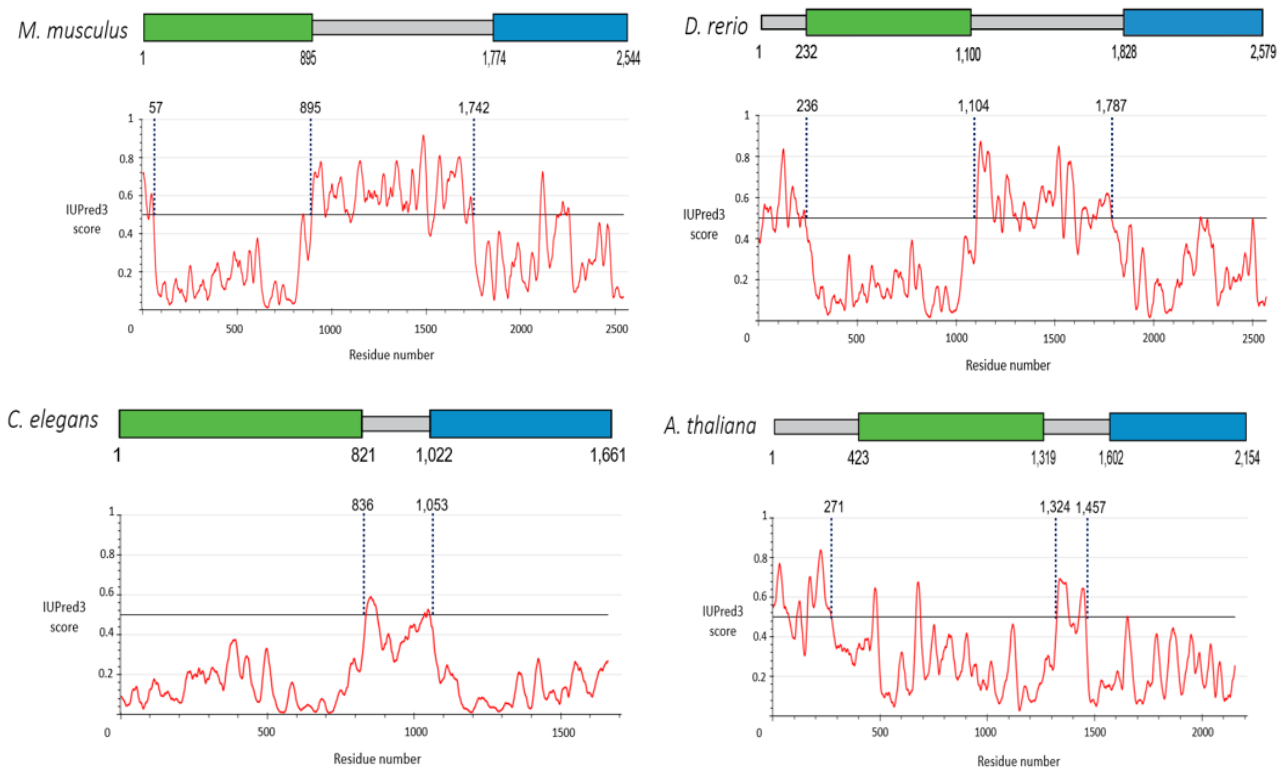


**Figure S3.** Pol θ linker domains are disordered. Protein disorder was predicted using IUPred3 long disorder and strong smoothing tools. Disordered residues have scores above 0.5 (horizontal line). Domain boundaries were predicted based on pairwise alignments with human helicase-like and polymerase domains.

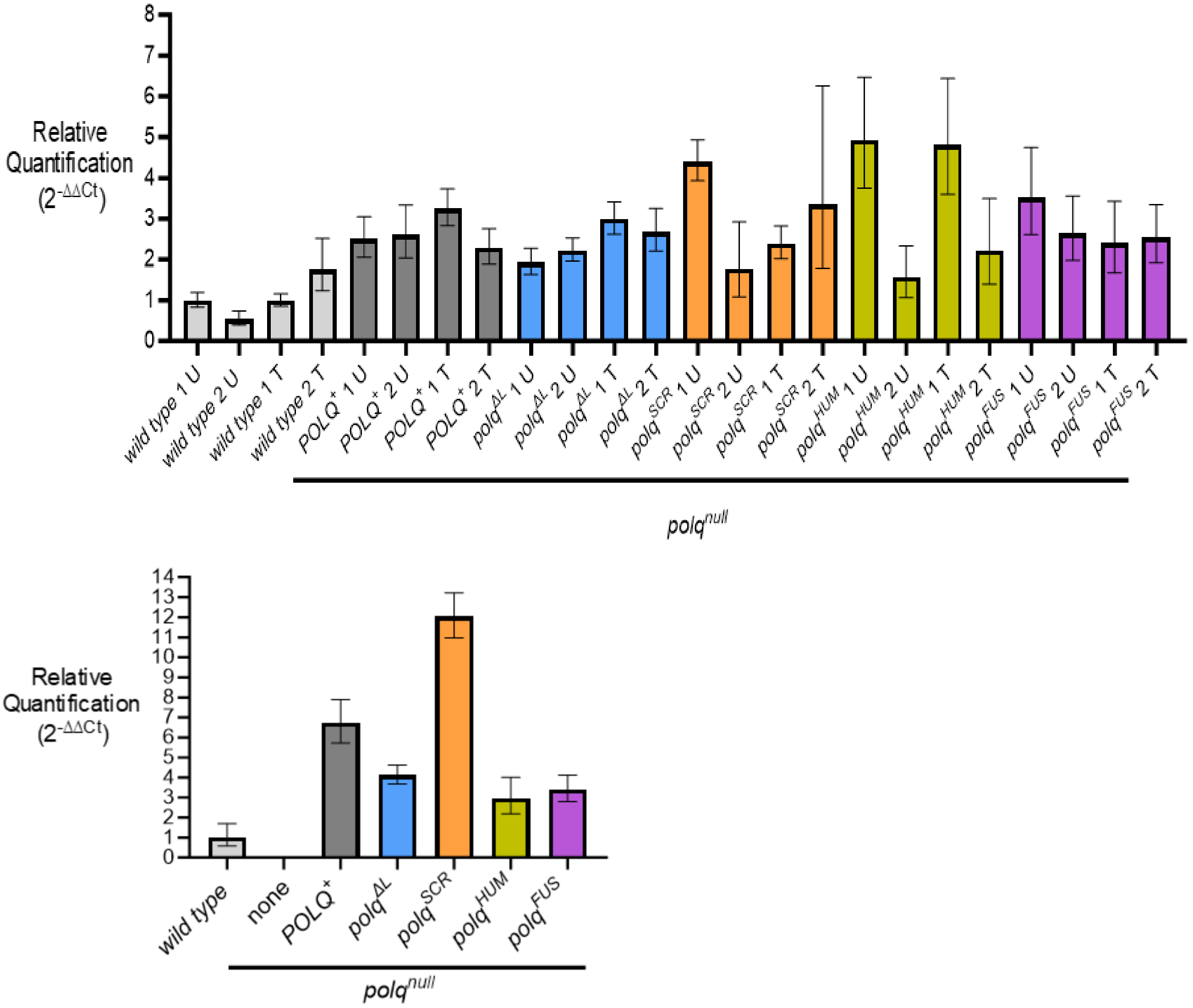


**Figure S4.** *polq*-mutant transgenes are expressed in larvae and adult flies. (**Upper panel**) *POLQ* mRNA expression in larvae homozygous for transgenes and $polq^{null}$ versus wild type. U=untreated. T=treated with 0.002% nitrogen mustard. RNA was extracted from larvae three days after treatment. 1 and 2 represent two separately-derived samples. Untreated and treated samples were analyzed in two separate qPCR experiments. Untreated or treated RQ values are shown relative to calibrators wild type 1U or 1T, respectively. Untreated RQ data are also shown in Figure 4B. (**Lower panel**) *POLQ* mRNA expression in adult flies. None=no transgene ($polq^{null}$). Calibrator=wild type. Both panels: endogenous control=*rp49* expression. Shown are RQmin and RQmax confidence intervals.

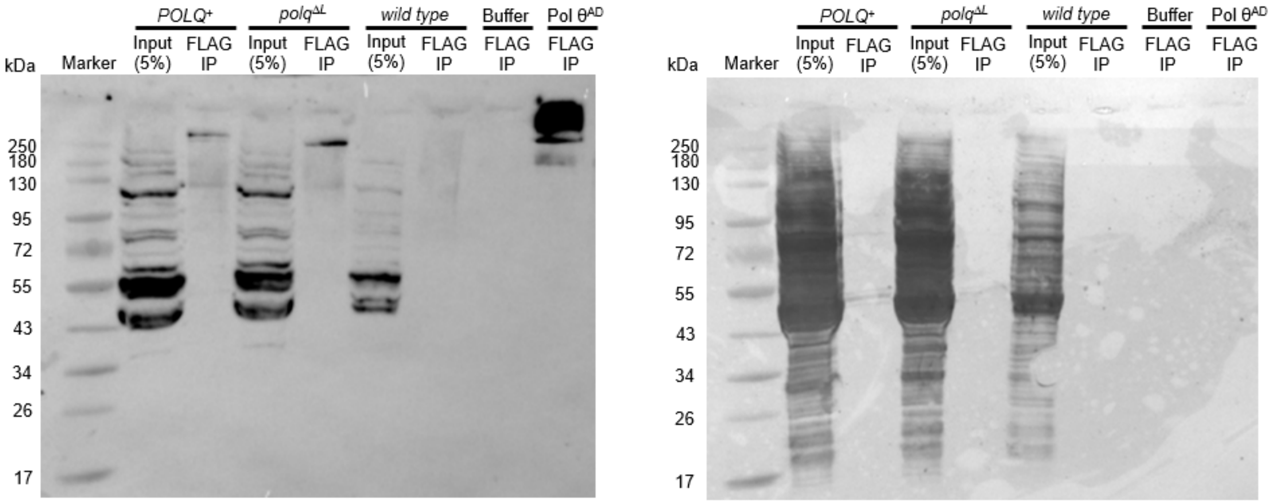


**Figure S5.** (**Left panel**) Uncropped blot image with overlaid marker shown in Figure 2C and additional controls. Buffer IP=anti-FLAG beads were incubated with lysis buffer only, Pol $\theta^{AD}$ IP=beads were incubated with 30 μg of FLAG-tagged ATPase-dead Pol θ described in Beagan *et al.* 2017. Proteins were detected using an anti-FLAG primary antibody. (**Right panel**) Reversible protein stain of membrane shown in left panel using G-Biosciences BLOT-FastStain.

A

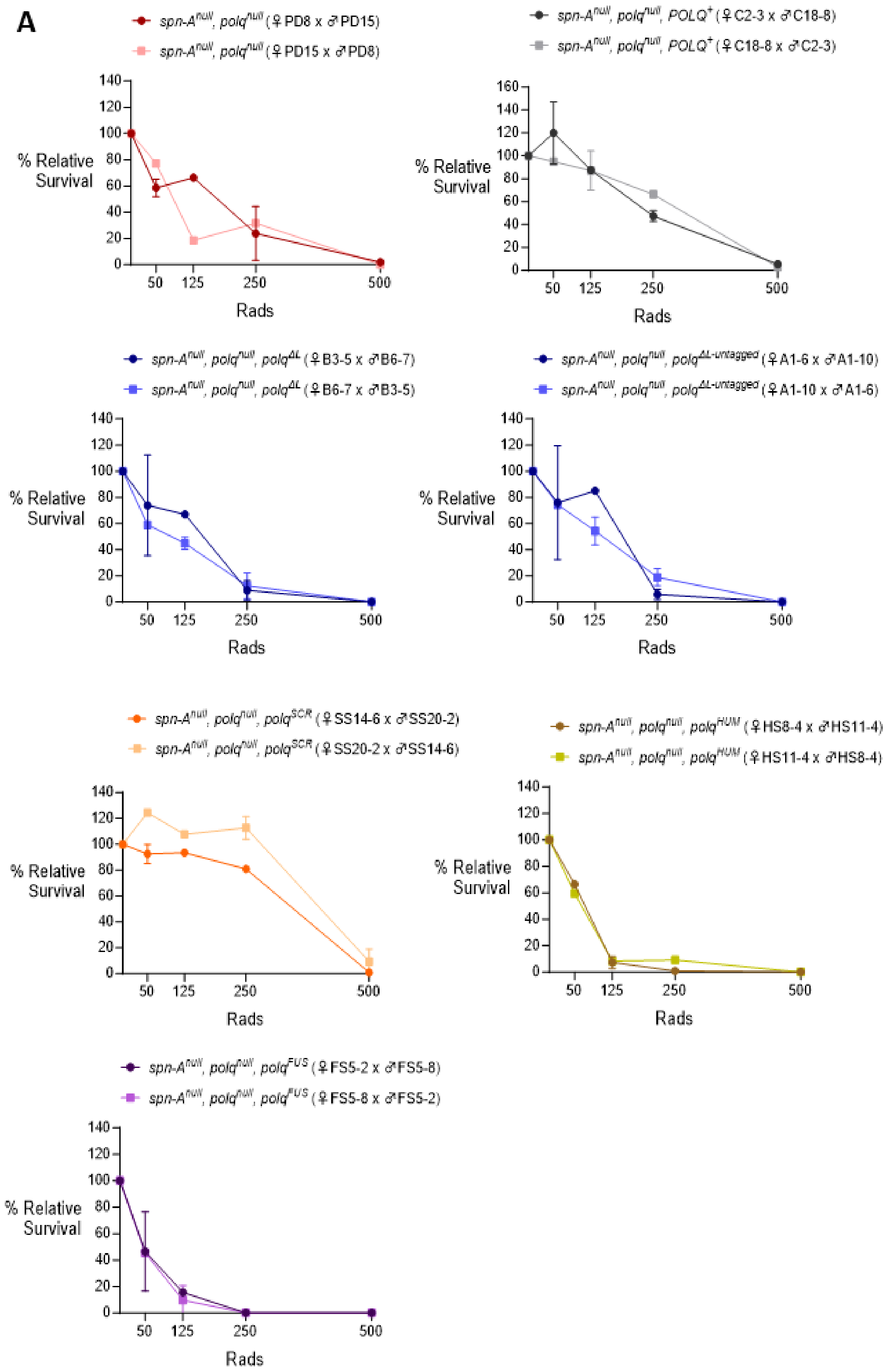
spn-A^null, polq^null (♀PD8 x ♂PD15)
spn-A^null, polq^null (♀PD15 x ♂PD8)
spn-A^null, polq^null, POLQ^+ (♀C2-3 x ♂C18-8)
spn-A^null, polq^null, POLQ^+ (♀C18-8 x ♂C2-3)
spn-A^null, polq^null, polq^ΔL (♀B3-5 x ♂B6-7)
spn-A^null, polq^null, polq^ΔL (♀B6-7 x ♂B3-5)
spn-A^null, polq^null, polq^ΔL-untagged (♀A1-6 x ♂A1-10)
spn-A^null, polq^null, polq^ΔL-untagged (♀A1-10 x ♂A1-6)
spn-A^null, polq^null, polq^SCR (♀SS14-6 x ♂SS20-2)
spn-A^null, polq^null, polq^SCR (♀SS20-2 x ♂SS14-6)
spn-A^null, polq^null, polq^HUM (♀HS8-4 x ♂HS11-4)
spn-A^null, polq^null, polq^HUM (♀HS11-4 x ♂HS8-4)
spn-A^null, polq^null, polq^FUS (♀FS5-2 x ♂FS5-8)
spn-A^null, polq^null, polq^FUS (♀FS5-8 x ♂FS5-2)
% Relative Survival
Rads
50
125
250
500
0
20
40
60
80
100
120
140
160

B

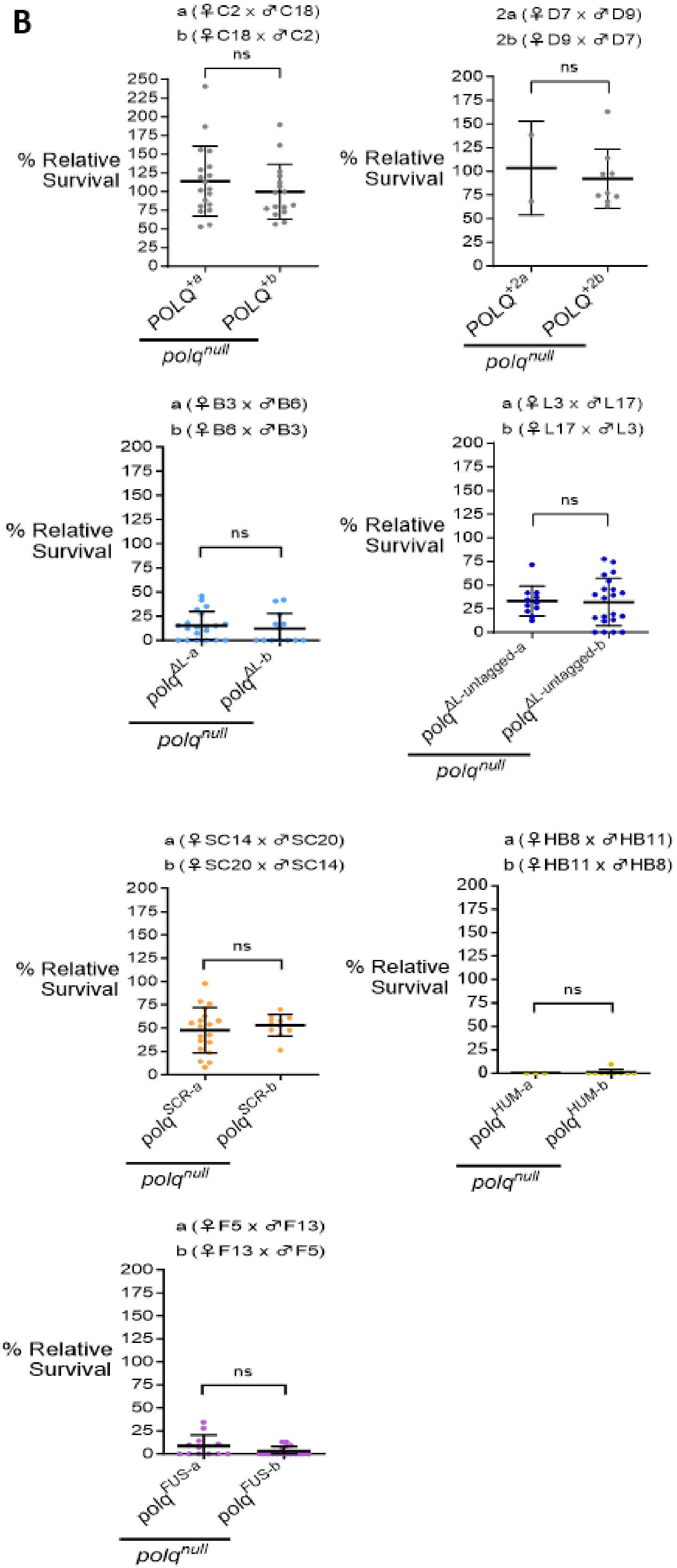
a (♀C2 x ♂C18)
b (♀C18 x ♂C2)
ns
% Relative Survival
POLQ+a
POLQ+b
polqnull
2a (♀D7 x ♂D9)
2b (♀D9 x ♂D7)
ns
% Relative Survival
POLQ+2a
POLQ+2b
polqnull
a (♀B3 x ♂B6)
b (♀B6 x ♂B3)
ns
% Relative Survival
polqΔL-a
polqΔL-b
polqnull
a (♀L3 x ♂L17)
b (♀L17 x ♂L3)
ns
% Relative Survival
polqΔL-untagged-a
polqΔL-untagged-b
polqnull
a (♀SC14 x ♂SC20)
b (♀SC20 x ♂SC14)
ns
% Relative Survival
polqSCR-a
polqSCR-b
polqnull
a (♀HB8 x ♂HB11)
b (♀HB11 x ♂HB8)
ns
% Relative Survival
polqHUM-a
polqHUM-b
polqnull
a (♀F5 x ♂F13)
b (♀F13 x ♂F5)
ns
% Relative Survival
polqFUS-a
polqFUS-b
polqnull

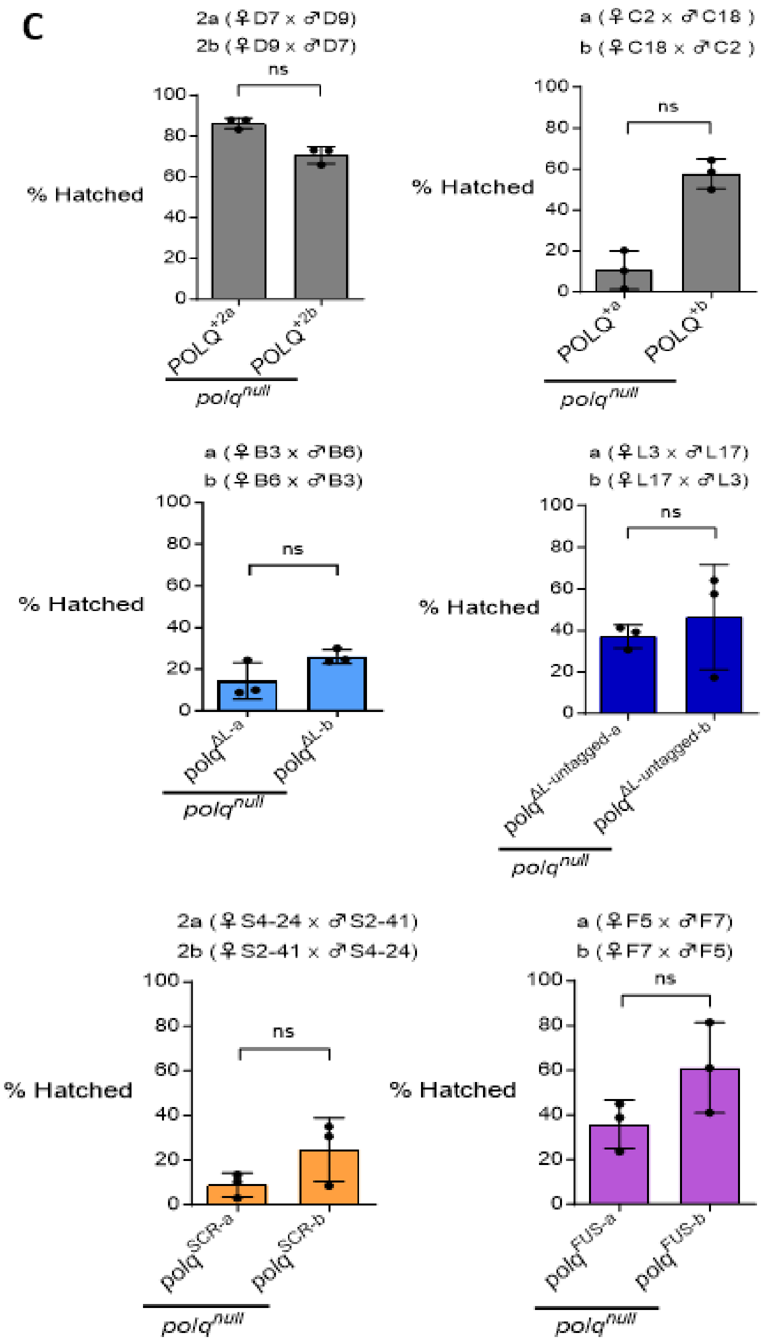


**Figure S6.** Survival comparisons of progeny from reciprocal crosses in IR, nitrogen mustard, and hatching assays. (**A**) Mean relative survival of irradiated larvae from reciprocal crosses in which male and female parent stocks were reversed. Shown are 1-2 replicates per reciprocal cross/dose and standard deviations. Statistical analyses are not shown since some crosses were performed once. (**B**) Mean relative survival of larvae treated with 0.007% nitrogen mustard. Shown are 2-21 replicates per reciprocal cross/dose and standard deviations, Statistical comparisons were made with Mann-Whitney tests. (**C**) Mean egg hatching rates. Parental flies were homozygous for the specified genotype. Shown are 3 replicates per reciprocal cross and standard deviations. Statistical comparisons were made with Mann-Whitney tests. Stock identifiers are listed in parentheses for all assays. Reciprocal cross data shown in each panel were combined for each genotype in Figure 3, Figure 5, and Supplementary Figure S7.

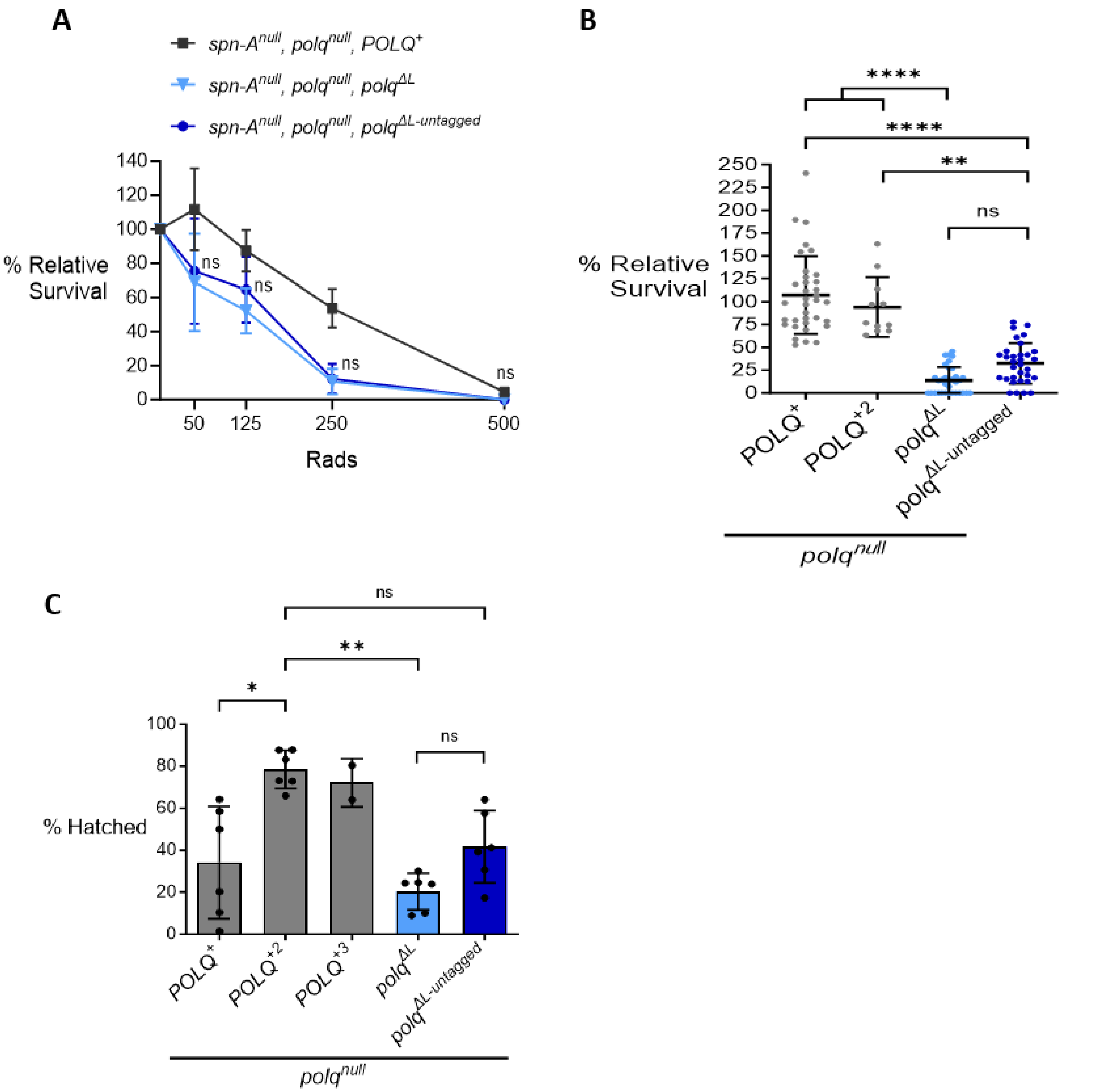


**Figure S7.** No significant differences between *polq*$^{\Delta L}$ and *polq*$^{\Delta L\text{-}untagged}$ mutants were observed in sensitivity and hatching assays. (**A**) Survival of irradiated larvae to adulthood relative to untreated larvae. The *polq*$^{\Delta L\text{-}untagged}$ transgene does not encode a N-terminal 3xFLAG tag. Each point represents mean relative survival of progeny from 3-4 crosses and standard deviations are shown. Comparisons between *spn-A*$^{null}$*, polq*$^{null}$*, polq*$^{\Delta L}$ and *spn-A*$^{null}$*, polq*$^{null}$*, polq*$^{\Delta L\text{-}untagged}$ were not significant (two-way ANOVA and Sidak's multiple comparisons test, all comparisons shown in Supplementary Table S6). (**B**) Relative survival of larvae treated with 0.007% nitrogen mustard. Shown are mean survival from 11-35 replicates per genotype and standard deviations. Comparisons were made using a Kruskal-Wallis ANOVA and Dunn's multiple comparisons test. (**C**) Mean egg hatching rates from 2-6 replicate crosses for each homozygous genotype. The *POLQ*$^{+2}$ cross included different parental stocks, but flies had the same genotype as *POLQ*$^{+}$. The *POLQ*$^{+3}$ cross had flies in which the *POLQ*$^{+}$ transgene was integrated at *ZH-68D2-attP* on chromosome 3 instead of *ZH-68E-attP*. Shown are standard deviations and comparisons using a Kruskal-Wallis ANOVA and Dunn's multiple comparisons test, unspecified comparisons were not significant. All panels: *p=0.01 to 0.05, **p=0.001 to 0.01, ****p=<0.0001, ns=not significant.

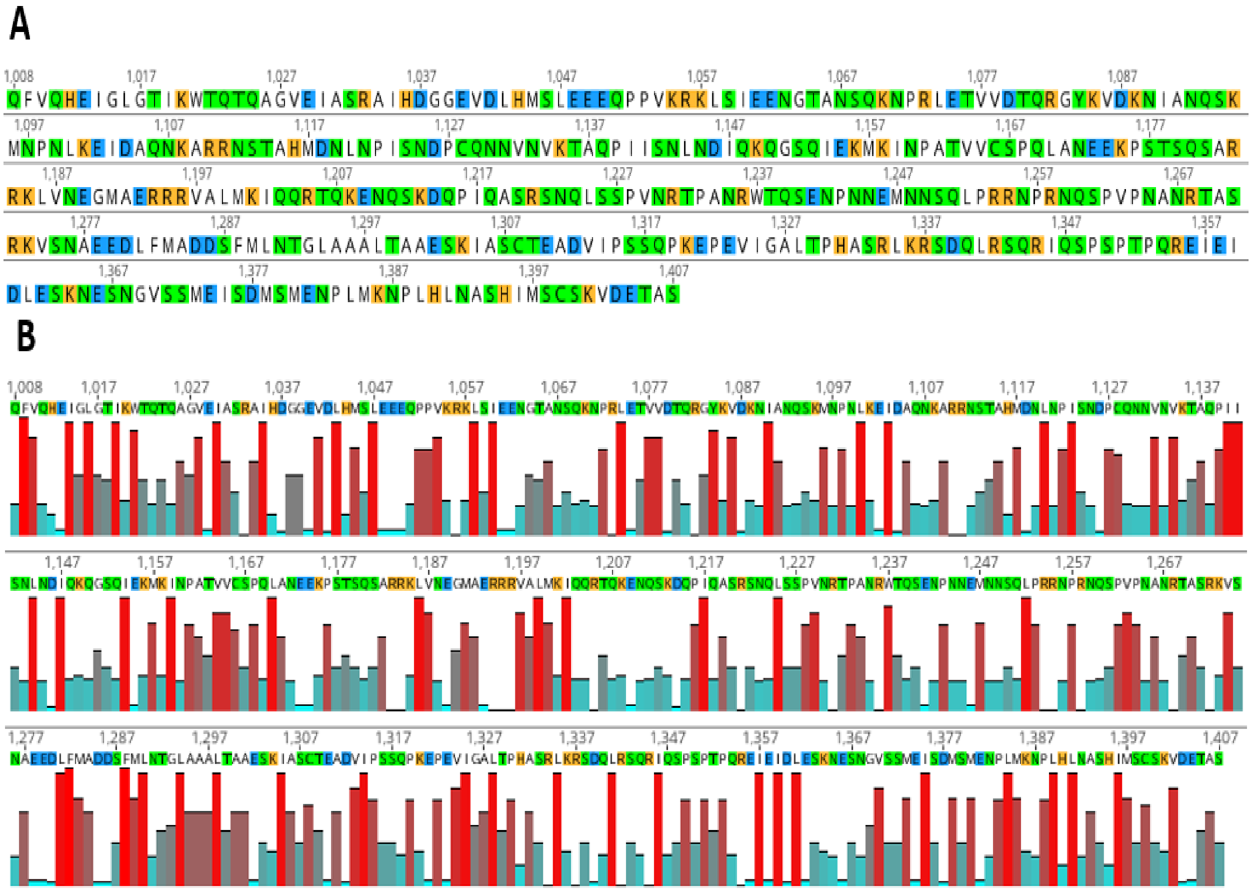

**Figure S8.** (**A**) Map of polar and charged residues within the *Drosophila melanogaster* Pol θ linker domain. Uncharged polar=green, negative=blue, positive=orange. (**B**) The *Drosophila melanogaster* Pol θ linker domain has clusters of hydrophilic residues. Hydrophobic=red bar, hydrophilic=blue bar. Panels were made in Geneious Prime.

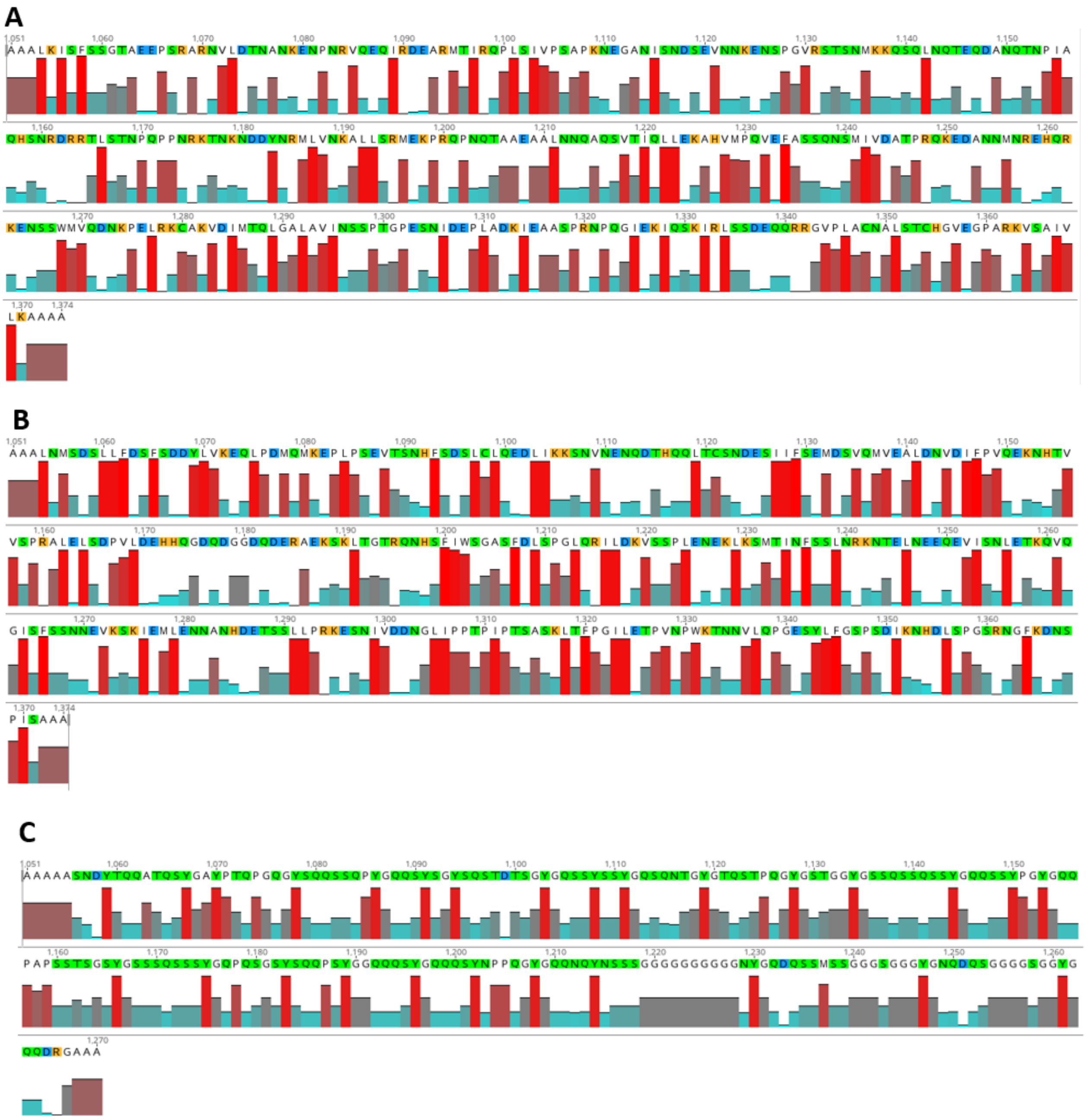


**Figure S9.** The scrambled (**A**), human (**B**), and FUS (**C**) linkers have clusters of hydrophilic residues and the FUS linker is mostly polar. Hydrophobic=red bar, hydrophilic=blue bar. Residues: uncharged polar=green, negative=blue, positive=orange. Panels were made in Geneious Prime.

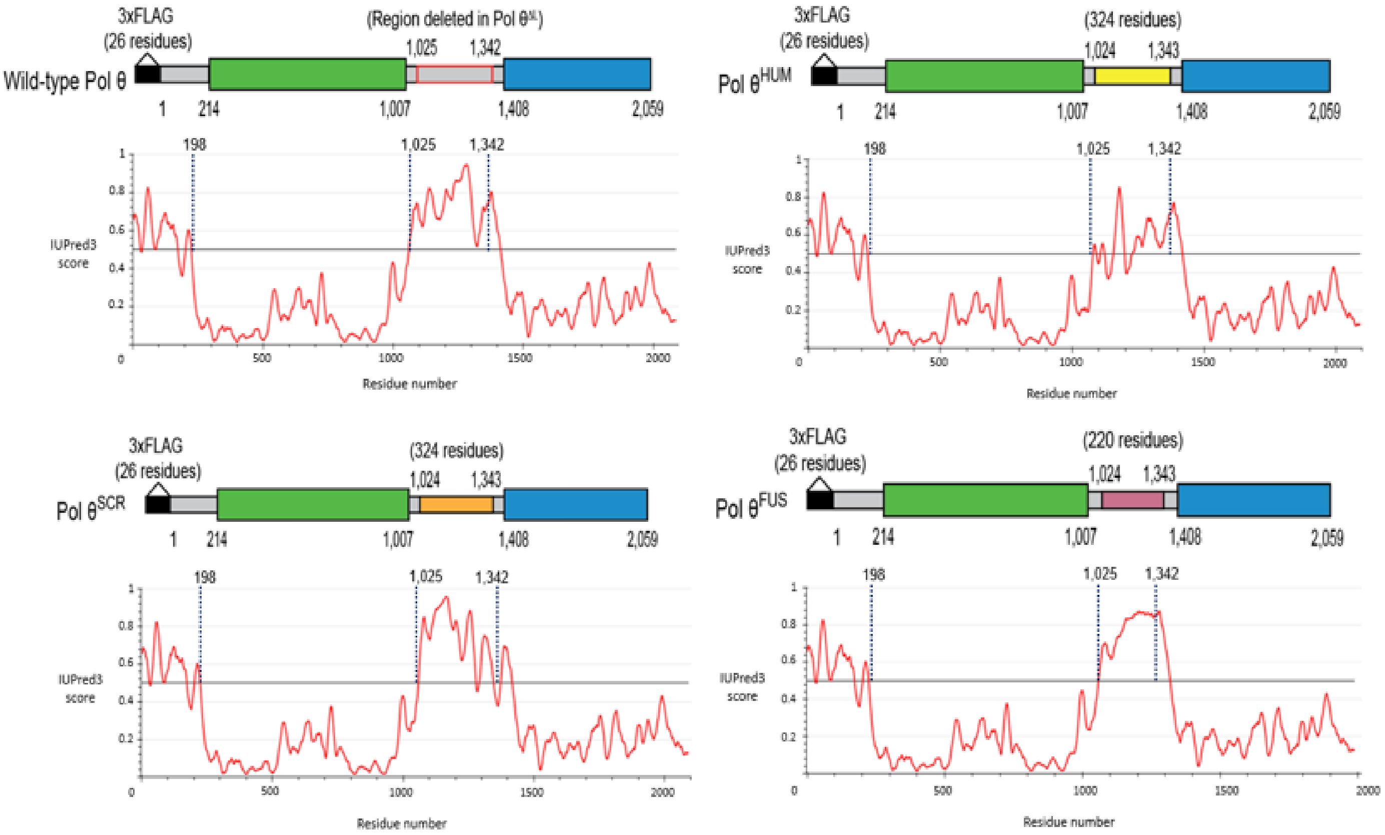


**Figure S10.** Replacement-linker sequences are disordered in *Drosophila melanogaster* Pol θ. Protein disorder was predicted using IUPred3 in the same way as other figures.

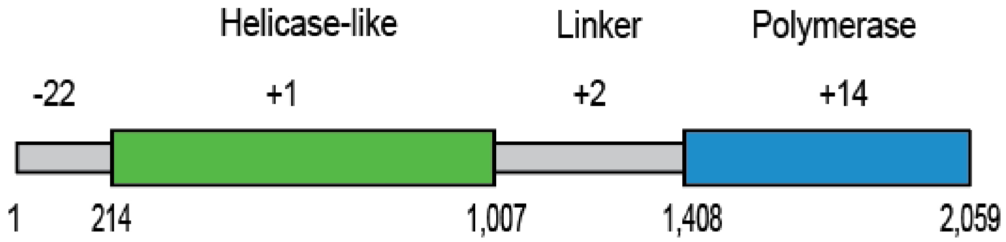


**Figure S11.** Overall charges of domains in *Drosophila melanogaster* Pol θ. Calculated at pH 7 in Geneious Prime. Residues 1-213= N-terminal undefined region.

**Table S1.** Primer sequences.

| Primer # | Sequence (5' to 3') |
|---|---|
| **1** | CCATCGTGGAGCAGTACGA |
| **2** | CTTCGTCTACCACCACCATGC |
| **3** | AAGCAACTGATGGCCATGG |
| **4** | CATGGCACTGTCCACAATCAGC |
| **5** | TTACAAGGATGACGATGACAAG |
| **6** | ACTGATTGGTGAAGCTCGAC |
| **7** | TTCCGGTGGCCACACACATC |
| **8** | ACAATAAACCAGAGCTGCGC |
| **9** | CCTTGTCCAAGATGCGTTGC |
| **10** | AGCAAAGTTACAATCCCCCG |
| **11** | CACTTGTAACAAGCAGTTTGC |
| **12** | AACGACTGCGAGAATGCCAT |
| **13** | ATCCGCCCAGCATACAGG |
| **14** | TCGATCCGTAACCGATGTTGG |

**Table S2.** gBlock sequences.

| gBlock | Sequence including NotI restriction sites and adapters (5' to 3') |
|---|---|
| Scrambled *Drosophila melanogaster* Pol θ linker | AGGACTACTCGATTCGTCGAACGTCGACTGAATGCGGCCGCTCT CAAGATCAGCTTTTCGTCGGGTACTGCCGAGGAACCCTCGAGGG CGCGGAACGTGCTGGATACTAACGCTAATAAGGAGAACCCCAAC CGGGTCCAGGAGCAAATTCGTGATGAGGCACGAATGACCATAAG GCAGCCGTTGAGTATAGTACCGAGTGCCCCTAAAAACGAAGGCG CTAATATCTCGAATGATAGTGAAGTAAATAACAAGGAGAACTCCCC TGGAGTACGTTCCACTAGTAATATGAAGAAGCAGTCGCAGTTGAA CCAGACGGAGCAAGATGCCAACCAGACGAATCCGATTGCGCAG CACTCCAACCGAGATCGAAGGACATTGTCCACGAACCCTCAGCC ACCCAACCGAAAGACAAATAAGAACGATGATTATAACCGGATGTT GGTTAACAAGGCCCTCTTGAGCCGTATGGAAAAGCCTCGGCAAC CGAACCAGACAGCTGCCGAAGCAGCCCTCAACAACCAGGCACA GTCCGTCACCATTCAACTCCTGGAGAAAGCGCACGTGATGCCGC AAGTTGAGTTCGCGAGCTCCCAGAATTCCATGATAGTTGATGCCA CACCACGACAGAAGGAAGACGCCAACAATATGAACAGGGAACAC CAGCGCAAGGAAAATTCCTCCTGGATGGTTCAAGACAATAAACCA GAGCTGCGCAAATGCGCCAAAGTAGACATTATGACGCAACTCGG AGCACTGGCAGTTATAAATTCCAGCCCGACTGGCCCTGAGTCGA ACATAGACGAGCCACTCGCAGATAAGATTGAGGCAGCTTCCCCT CGAAATCCGCAGGGTATAGAAAAAATCCAAAGTAAAATCCGGCTG AGTAGCGATGAGCAACAACGTCGCGGAGTACCTCTCGCCTGCAA TGCCTTGAGTACCTGTCATGGTGTTGAGGGTCCAGCACGGAAGG TTAGCGCGATTGTGCTCAAAGCTGCGGCCGCATACTGACCACGA TTGCCGTCACACGTTACTAGG |

| | |
|---|---|
| *Homo sapiens* Pol θ linker | AGGACTACTCGATTCGTCGAACGTCGACTGAATGCGGCCGCTCT<br>GAATATGAGTGATTCCCTCTTGTTCGATTCCTTTAGCGATGATTAT<br>CTGGTTAAAGAGCAATTGCCAGACATGCAAATGAAGGAGCCACT<br>GCCTAGTGAGGTGACCTCGAACCATTTCTCCGATAGTCTCTGCTT<br>GCAAGAGGATTTGATAAAGAAGAGCAATGTAAACGAAAATCAAGA<br>CACACACCAACAATTGACATGCAGCAACGATGAAAGTATTATATTC<br>TCGGAAATGGACAGCGTCCAGATGGTTGAGGCGTTGGATAACGT<br>GGACATTTTCCCGGTACAGGAGAAGAATCATACAGTGGTAAGCCC<br>CCGCGCTTTGGAGCTGTCCGACCCCGTTTTGGACGAGCATCATC<br>AAGGCGACCAAGATGGAGGAGACCAGGACGAACGCGCGGAAAA<br>ATCCAAGCTGACAGGAACCCGTCAGAACCACTCGTTCATCTGGA<br>GTGGAGCCTCCTTCGACCTCAGCCCGGGTTTGCAACGCATCTTG<br>GACAAGGTAAGTTCCCCCCTGGAGAATGAGAAATTGAAATCGATG<br>ACGATTAACTTTTCCTCGCTCAATCGCAAAAACACGGAATTGAAT<br>GAAGAACAGGAAGTCATCAGTAACCTCGAGACAAAACAGGTGCA<br>AGGCATCTCCTTTTCCTCCAACAATGAAGTAAAATCCAAAATCGA<br>GATGCTGGAGAATAACGCGAATCATGATGAAACAAGCTCGCTGCT<br>CCCACGAAAAGAGTCGAACATAGTCGATGACAATGGATTGATCCC<br>TCCAACGCCCATTCCAACGAGTGCTTCGAAGTTGACCTTCCCTG<br>GTATCTTGGAGACTCCCGTGAATCCGTGGAAGACAAATAATGTCC<br>TCCAGCCCGGAGAGAGTTACCTGTTCGGATCGCCTAGCGACATC<br>AAGAATCATGATCTGTCCCCCGGTTCGCGCAATGGTTTTAAAGAT<br>AACTCCCCTATCAGTGCGGCCGCATACTGACCACGATTGCCGTC<br>ACACGTTACTAGG |
| FUS linker | GACTGAATGCGGCCGCTGCTGCCAGTAATGATTATACACAACAAG<br>CGACTCAGTCGTATGGTGCGTATCCGACTCAACCCGGCCAGGGA<br>TACTCGCAGCAGTCCAGTCAACCTTACGGCCAACAGAGCTACAG<br>CGGTTATTCCCAATCGACAGATACATCGGGCTATGGACAGAGTAG<br>CTACAGCAGTTACGGTCAAAGTCAAAACACTGGTTATGGCACACA<br>GTCCACGCCTCAGGGCTACGGTAGCACAGGTGGTTACGGAAGTA<br>GTCAGTCCAGCCAAAGCTCCTATGGTCAGCAGAGCAGTTATCCC<br>GGTTATGGCCAGCAGCCTGCCCCAAGCAGTACCTCCGGCAGCTA<br>CGGCTCCTCCTCGCAATCGTCCAGTTACGGACAACCTCAAAGTG<br>GTAGCTACAGTCAGCAACCCTCGTATGGCGGCCAACAGCAGAGC<br>TATGGTCAGCAGCAAAGTTACAATCCCCCGCAAGGATATGGTCAA<br>CAGAATCAGTATAACAGTAGTTCGGGCGGAGGCGGAGGAGGTG<br>GTGGTGGCGGCAATTATGGACAAGATCAGAGTTCGATGAGTTCC<br>GGAGGCGGCAGTGGCGGCGGCTACGGAAATCAGGACCAGAGC<br>GGTGGAGGCGGATCGGGAGGTTACGGACAGCAGGATCGAGGAG<br>CGGCCGCATACTGAAT |

**Table S3.** No-RT control Ct values of RNA used for cDNA preparation.

| Category | Well | Sample | Ct |
|---|---|---|---|
| U | A1 | *POLQ*$^{+}$ 1 | undetermined |
| U | A2 | *POLQ*$^{+}$ 1 | 37.437 |
| U | A3 | *POLQ*$^{+}$ 1 | 38.808 |
| U | A4 | *POLQ*$^{+}$ 2 | undetermined |
| U | A5 | *POLQ*$^{+}$ 2 | undetermined |
| U | A6 | *POLQ*$^{+}$ 2 | undetermined |
| U | A7 | *polq*$^{\Delta L}$ *1* | 37.079 |
| U | A8 | *polq*$^{\Delta L}$ *1* | 36.096 |
| U | A9 | *polq*$^{\Delta L}$ *1* | 36.323 |
| U | A10 | *polq*$^{\Delta L}$ 2 | undetermined |
| U | A11 | *polq*$^{\Delta L}$ 2 | undetermined |
| U | A12 | *polq*$^{\Delta L}$ 2 | 38.949 |
| U | B1 | *polq*$^{SCR}$ *1* | undetermined |
| U | B2 | *polq*$^{SCR}$ *1* | 37.031 |
| U | B3 | *polq*$^{SCR}$ *1* | undetermined |
| U | B4 | *polq*$^{SCR}$ 2 | 38.716 |
| U | B5 | *polq*$^{SCR}$ 2 | undetermined |
| U | B6 | *polq*$^{SCR}$ 2 | undetermined |
| U | B7 | *polq*$^{HUM}$ *1* | 39.550 |
| U | B8 | *polq*$^{HUM}$ *1* | undetermined |
| U | B9 | *polq*$^{HUM}$ *1* | 38.258 |
| U | B10 | *polq*$^{HUM}$ 2 | 38.254 |
| U | B11 | *polq*$^{HUM}$ 2 | 39.363 |
| U | B12 | *polq*$^{HUM}$ 2 | undetermined |
| U | C1 | *polq*$^{FUS}$ 1 | 37.127 |
| U | C2 | *polq*$^{FUS}$ 1 | 38.021 |
| U | C3 | *polq*$^{FUS}$ 1 | 38.097 |
| U | C4 | *polq*$^{FUS}$ 2 | 37.899 |
| U | C5 | *polq*$^{FUS}$ 2 | 37.196 |
| U | C6 | *polq*$^{FUS}$ 2 | 36.482 |
| U | C7 | *wild type* 1 | 36.554 |
| U | C8 | *wild type* 1 | 38.916 |
| U | C9 | *wild type* 1 | 37.823 |
| U | C10 | *wild type* 2 | undetermined |
| U | C11 | *wild type* 2 | 38.405 |
| U | C12 | *wild type* 2 | undetermined |
| U | D1 | No template control | undetermined |
| U | D2 | No template control | undetermined |
| U | D3 | No template control | undetermined |
| U | D4 | *POLQ*$^{+}$ 1 positive control | 29.776 |
| T | A1 | *POLQ*$^{+}$ 1 | undetermined |
| T | A2 | *POLQ*$^{+}$ 1 | 38.097 |
| T | A3 | *POLQ*$^{+}$ 1 | undetermined |
| T | A4 | *POLQ*$^{+}$ 2 | undetermined |
| T | A5 | *POLQ*$^{+}$ 2 | undetermined |
| T | A6 | *POLQ*$^{+}$ 2 | undetermined |
| T | A7 | *polq*$^{\Delta L}$ *1* | 38.034 |
| T | A8 | *polq*$^{\Delta L}$ *1* | undetermined |
| T | A9 | *polq*$^{\Delta L}$ *1* | undetermined |

| | | | |
|---|---|---|---|
| T | A10 | *polq*$^{\Delta L}$ *2* | undetermined |
| T | A11 | *polq*$^{\Delta L}$ *2* | 39.822 |
| T | A12 | *polq*$^{\Delta L}$ *2* | undetermined |
| T | B1 | *polq*$^{SCR}$ *1* | 39.787 |
| T | B2 | *polq*$^{SCR}$ *1* | undetermined |
| T | B3 | *polq*$^{SCR}$ *1* | undetermined |
| T | B4 | *polq*$^{SCR}$ 2 | undetermined |
| T | B5 | *polq*$^{SCR}$ 2 | undetermined |
| T | B6 | *polq*$^{SCR}$ 2 | 38.319 |
| T | B7 | *polq*$^{HUM}$ *1* | 39.805 |
| T | B8 | *polq*$^{HUM}$ *1* | undetermined |
| T | B9 | *polq*$^{HUM}$ *1* | 39.194 |
| T | B10 | *polq*$^{HUM}$ 2 | 37.690 |
| T | B11 | *polq*$^{HUM}$ 2 | undetermined |
| T | B12 | *polq*$^{HUM}$ 2 | undetermined |
| T | C1 | *polq*$^{FUS}$ 1 | undetermined |
| T | C2 | *polq*$^{FUS}$ 1 | undetermined |
| T | C3 | *polq*$^{FUS}$ 1 | undetermined |
| T | C4 | *polq*$^{FUS}$ 2 | undetermined |
| T | C5 | *polq*$^{FUS}$ 2 | undetermined |
| T | C6 | *polq*$^{FUS}$ 2 | undetermined |
| T | C7 | *wild type* 1 | 38.222 |
| T | C8 | *wild type* 1 | 39.277 |
| T | C9 | *wild type* 1 | undetermined |
| T | C10 | *wild type* 2 | undetermined |
| T | C11 | *wild type* 2 | undetermined |
| T | C12 | *wild type* 2 | undetermined |
| T | D1 | No template control | undetermined |
| T | D2 | No template control | undetermined |
| T | D3 | No template control | undetermined |
| T | D4 | *POLQ*$^{+}$ 1 positive control | 30.204 |
| A | F1 | *POLQ*$^{+}$ | undetermined |
| A | F2 | *POLQ*$^{+}$ | 39.566 |
| A | F3 | *POLQ*$^{+}$ | 39.536 |
| A | F4 | *polq*$^{\Delta L}$ | 36.951 |
| A | F5 | *polq*$^{\Delta L}$ | 38.229 |
| A | F6 | *polq*$^{\Delta L}$ | 35.850 |
| A | F7 | *polq*$^{SCR}$ | 38.138 |
| A | F8 | *polq*$^{SCR}$ | undetermined |
| A | F9 | *polq*$^{SCR}$ | 39.987 |
| A | F10 | *polq*$^{HUM}$ | 38.046 |
| A | F11 | *polq*$^{HUM}$ | undetermined |
| A | F12 | *polq*$^{HUM}$ | 37.821 |
| A | G1 | *polq*$^{FUS}$ | 39.415 |
| A | G2 | *polq*$^{FUS}$ | 38.857 |
| A | G3 | *polq*$^{FUS}$ | undetermined |
| A | G4 | *wild type* | 39.517 |
| A | G5 | *wild type* | 37.921 |
| A | G6 | *wild type* | undetermined |
| A | G7 | *polq*$^{null}$ | undetermined |
| A | G8 | *polq*$^{null}$ | undetermined |
| A | G9 | *polq*$^{null}$ | undetermined |
| A | D4 | No template control | 39.126 |
| A | D5 | No template control | undetermined |

| | | | |
|---|---|---|---|
| A | D6 | No template control | undetermined |
| A | A1 | *POLQ*$^{+}$ positive control | 26.778 |

*POLQ* amplification. U=untreated larvae, T=larvae treated with 0.002% nitrogen mustard, A=untreated adult flies. A positive control was included for each category to show each master mix could amplify cDNA. Undetermined=fluorescence did not exceed the Ct threshold.

**Table S4.** Protein sequence accession numbers.

| Protein | Organism | Database | Accession number |
|---|---|---|---|
| Pol θ | *Drosophila melanogaster* | UniProt | O18475 |
| Pol θ | *Homo sapiens* | UniProt | O75417-1 |
| Pol θ | *Bos taurus* | UniProt | E1BMC0 |
| Pol θ | *Rattus norvegicus* | UniProt | D4A628 |
| Pol θ | *Mus musculus* | UniProt | Q8CGS6-1 |
| Pol θ | *Tursiops truncatus* | UniProt | A0A6J3RCQ3 |
| Pol θ | *Gallus gallus* | UniProt | A0A8V1A4V3 |
| Pol θ | *Xenopus laevis* | UniProt | A0A977PDH1 |
| Pol θ | *Danio rerio* | UniProt | A0A8M9Q371 |
| Pol θ | *Caenorhabditis elegans* | UniProt | A0FLQ6 |
| Pol θ | *Arabidopsis thaliana* | NCBI | BAD93700.1 |
| Pol θ | *Drosophila erecta* | NCBI | XP_001980357 |
| Pol θ | *Drosophila obscura* | NCBI | XP_022234266 |
| Pol θ | *Drosophila hydei* | NCBI | XP_023168898 |
| Pol θ | *Drosophila persimilis* | NCBI | XP_026843212 |
| Pol θ | *Drosophila serrata* | NCBI | XP_020811369 |
| FUS | *Homo sapiens* | UniProt | P35637-1 |

**Table S5.** Comprehensive significance summary of IR sensitivity data in Figure 3A.

| IR Dose (rads) | Comparison | Summary | P value |
|---|---|---|---|
| 50 | *spn-A*$^{null}$ **vs.** *spn-A*$^{null}$, *polq*$^{null}$ | ns | 0.0630 |
| 50 | *spn-A*$^{null}$ **vs.** *spn-A*$^{null}$, *polq*$^{null}$, *polq*$^{\Delta L}$ | ns | 0.1438 |
| 50 | *spn-A*$^{null}$ **vs.** *spn-A*$^{null}$, *polq*$^{null}$, *POLQ*$^{+}$ | ns | 0.7628 |
| 50 | *spn-A*$^{null}$, *polq*$^{null}$ **vs.** *spn-A*$^{null}$, *polq*$^{null}$, *polq*$^{\Delta L}$ | ns | 0.9996 |
| 50 | *spn-A*$^{null}$, *polq*$^{null}$ **vs.** *spn-A*$^{null}$, *polq*$^{null}$, *POLQ*$^{+}$ | ** | 0.0018 |
| 50 | *spn-A*$^{null}$, *polq*$^{null}$, *polq*$^{\Delta L}$ **vs.** *spn-A*$^{null}$, *polq*$^{null}$, *POLQ*$^{+}$ | ** | 0.0052 |
| 125 | *spn-A*$^{null}$ **vs.** *spn-A*$^{null}$, *polq*$^{null}$ | ** | 0.0035 |
| 125 | *spn-A*$^{null}$ **vs.** *spn-A*$^{null}$, *polq*$^{null}$, *polq*$^{\Delta L}$ | ns | 0.1716 |
| 125 | *spn-A*$^{null}$ **vs.** *spn-A*$^{null}$, *polq*$^{null}$, *POLQ*$^{+}$ | ns | 0.9815 |
| 125 | *spn-A*$^{null}$, *polq*$^{null}$ **vs.** *spn-A*$^{null}$, *polq*$^{null}$, *polq*$^{\Delta L}$ | ns | 0.6087 |
| 125 | *spn-A*$^{null}$, *polq*$^{null}$ **vs.** *spn-A*$^{null}$, *polq*$^{null}$, *POLQ*$^{+}$ | *** | 0.0004 |
| 125 | *spn-A*$^{null}$, *polq*$^{null}$, *polq*$^{\Delta L}$ **vs.** *spn-A*$^{null}$, *polq*$^{null}$, *POLQ*$^{+}$ | * | 0.0316 |
| 250 | *spn-A*$^{null}$ **vs.** *spn-A*$^{null}$, *polq*$^{null}$ | ** | 0.0035 |
| 250 | *spn-A*$^{null}$ **vs.** *spn-A*$^{null}$, *polq*$^{null}$, *polq*$^{\Delta L}$ | **** | <0.0001 |
| 250 | *spn-A*$^{null}$ **vs.** *spn-A*$^{null}$, *polq*$^{null}$, *POLQ*$^{+}$ | ns | 0.6592 |
| 250 | *spn-A*$^{null}$, *polq*$^{null}$ **vs.** *spn-A*$^{null}$, *polq*$^{null}$, *polq*$^{\Delta L}$ | ns | 0.6694 |
| 250 | *spn-A*$^{null}$, *polq*$^{null}$ **vs.** *spn-A*$^{null}$, *polq*$^{null}$, *POLQ*$^{+}$ | ns | 0.1496 |
| 250 | *spn-A*$^{null}$, *polq*$^{null}$, *polq*$^{\Delta L}$ **vs.** *spn-A*$^{null}$, *polq*$^{null}$, *POLQ*$^{+}$ | ** | 0.0023 |
| 500 | *spn-A*$^{null}$ **vs.** *spn-A*$^{null}$, *polq*$^{null}$ | ** | 0.0012 |
| 500 | *spn-A*$^{null}$ **vs.** *spn-A*$^{null}$, *polq*$^{null}$, *polq*$^{\Delta L}$ | *** | 0.0008 |
| 500 | *spn-A*$^{null}$ **vs.** *spn-A*$^{null}$, *polq*$^{null}$, *POLQ*$^{+}$ | ** | 0.0027 |
| 500 | *spn-A*$^{null}$, *polq*$^{null}$ **vs.** *spn-A*$^{null}$, *polq*$^{null}$, *polq*$^{\Delta L}$ | ns | >0.9999 |
| 500 | *spn-A*$^{null}$, *polq*$^{null}$ **vs.** *spn-A*$^{null}$, *polq*$^{null}$, *POLQ*$^{+}$ | ns | >0.9999 |
| 500 | *spn-A*$^{null}$, *polq*$^{null}$, *polq*$^{\Delta L}$ **vs.** *spn-A*$^{null}$, *polq*$^{null}$, *POLQ*$^{+}$ | ns | 0.9994 |

Two-way ANOVA and Sidak's multiple comparisons test, *p=0.01 to 0.05, **p=0.001 to 0.01, ***p=0.0001 to 0.001, ****p=<0.0001, ns=not significant.

**Table S6.** Comprehensive significance summary of data in Supplementary Figure S7A.

| IR Dose (rads) | Comparison | Summary | P value |
|---|---|---|---|
| 50 | *spn-A$^{null}$, polq$^{null}$, polq$^{\Delta L\text{-}untagged}$* **vs.** *spn-A$^{null}$, polq$^{null}$, polq$^{\Delta L}$* | ns | 0.9272 |
| 50 | *spn-A$^{null}$, polq$^{null}$, polq$^{\Delta L\text{-}untagged}$* **vs.** *spn-A$^{null}$, polq$^{null}$, POLQ$^{+}$* | * | 0.0115 |
| 50 | *spn-A$^{null}$, polq$^{null}$, polq$^{\Delta L}$* **vs.** *spn-A$^{null}$, polq$^{null}$, POLQ$^{+}$* | ** | 0.0026 |
| 125 | *spn-A$^{null}$, polq$^{null}$, polq$^{\Delta L\text{-}untagged}$* **vs.** *spn-A$^{null}$, polq$^{null}$, polq$^{\Delta L}$* | ns | 0.6545 |
| 125 | *spn-A$^{null}$, polq$^{null}$, polq$^{\Delta L\text{-}untagged}$* **vs.** *spn-A$^{null}$, polq$^{null}$, POLQ$^{+}$* | ns | 0.1688 |
| 125 | *spn-A$^{null}$, polq$^{null}$, polq$^{\Delta L}$* **vs.** *spn-A$^{null}$, polq$^{null}$, POLQ$^{+}$* | * | 0.0150 |
| 250 | *spn-A$^{null}$, polq$^{null}$, polq$^{\Delta L\text{-}untagged}$* **vs.** *spn-A$^{null}$, polq$^{null}$, polq$^{\Delta L}$* | ns | 0.9981 |
| 250 | *spn-A$^{null}$, polq$^{null}$, polq$^{\Delta L\text{-}untagged}$* **vs.** *spn-A$^{null}$, polq$^{null}$, POLQ$^{+}$* | ** | 0.0018 |
| 250 | *spn-A$^{null}$, polq$^{null}$, polq$^{\Delta L}$* **vs.** *spn-A$^{null}$, polq$^{null}$, POLQ$^{+}$* | ** | 0.0012 |
| 500 | *spn-A$^{null}$, polq$^{null}$, polq$^{\Delta L\text{-}untagged}$* **vs.** *spn-A$^{null}$, polq$^{null}$, polq$^{\Delta L}$* | ns | >0.9999 |
| 500 | *spn-A$^{null}$, polq$^{null}$, polq$^{\Delta L\text{-}untagged}$* **vs.** *spn-A$^{null}$, polq$^{null}$, POLQ$^{+}$* | ns | 0.9733 |
| 500 | *spn-A$^{null}$, polq$^{null}$, polq$^{\Delta L}$* **vs.** *spn-A$^{null}$, polq$^{null}$, POLQ$^{+}$* | ns | 0.9733 |

Two-way ANOVA and Sidak's multiple comparisons test, *p=0.01 to 0.05, **p=0.001 to 0.01, ns=not significant.

**Table S7.** Comprehensive significance summary of comparisons in Figure 5A.

| Graph | IR Dose (rads) | Comparison | Summary | P value |
|---|---|---|---|---|
| SCR | 50 | *spn-A$^{null}$, polq$^{null}$, polq$^{\Delta L}$* **vs.** *spn-A$^{null}$, polq$^{null}$, POLQ$^{+}$* | ** | 0.0015 |
| SCR | 50 | *spn-A$^{null}$, polq$^{null}$, polq$^{\Delta L}$* **vs.** *spn-A$^{null}$, polq$^{null}$, polq$^{SCR}$* | * | 0.0118 |
| SCR | 50 | *spn-A$^{null}$, polq$^{null}$, POLQ$^{+}$* **vs.** *spn-A$^{null}$, polq$^{null}$, polq$^{SCR}$* | ns | 0.8302 |
| SCR | 125 | *spn-A$^{null}$, polq$^{null}$, polq$^{\Delta L}$* **vs.** *spn-A$^{null}$, polq$^{null}$, POLQ$^{+}$* | ** | 0.0098 |
| SCR | 125 | *spn-A$^{null}$, polq$^{null}$, polq$^{\Delta L}$* **vs.** *spn-A$^{null}$, polq$^{null}$, polq$^{SCR}$* | *** | 0.0007 |
| SCR | 125 | *spn-A$^{null}$, polq$^{null}$, POLQ$^{+}$* **vs.** *spn-A$^{null}$, polq$^{null}$, polq$^{SCR}$* | ns | 0.6957 |
| SCR | 250 | *spn-A$^{null}$, polq$^{null}$, polq$^{\Delta L}$* **vs.** *spn-A$^{null}$, polq$^{null}$, POLQ$^{+}$* | *** | 0.0006 |
| SCR | 250 | *spn-A$^{null}$, polq$^{null}$, polq$^{\Delta L}$* **vs.** *spn-A$^{null}$, polq$^{null}$, polq$^{SCR}$* | **** | <0.0001 |
| SCR | 250 | *spn-A$^{null}$, polq$^{null}$, POLQ$^{+}$* **vs.** *spn-A$^{null}$, polq$^{null}$, polq$^{SCR}$* | *** | 0.0004 |
| SCR | 500 | *spn-A$^{null}$, polq$^{null}$, polq$^{\Delta L}$* **vs.** *spn-A$^{null}$, polq$^{null}$, POLQ$^{+}$* | ns | 0.9685 |
| SCR | 500 | *spn-A$^{null}$, polq$^{null}$, polq$^{\Delta L}$* **vs.** *spn-A$^{null}$, polq$^{null}$, polq$^{SCR}$* | ns | 0.9161 |
| SCR | 500 | *spn-A$^{null}$, polq$^{null}$, POLQ$^{+}$* **vs.** *spn-A$^{null}$, polq$^{null}$, polq$^{SCR}$* | ns | 0.9974 |
| HUM | 50 | *spn-A$^{null}$, polq$^{null}$, polq$^{\Delta L}$* **vs.** *spn-A$^{null}$, polq$^{null}$, POLQ$^{+}$* | *** | 0.0002 |
| HUM | 50 | *spn-A$^{null}$, polq$^{null}$, polq$^{\Delta L}$* **vs.** *spn-A$^{null}$, polq$^{null}$, polq$^{HUM}$* | ns | 0.9389 |
| HUM | 50 | *spn-A$^{null}$, polq$^{null}$, POLQ$^{+}$* **vs.** *spn-A$^{null}$, polq$^{null}$, polq$^{HUM}$* | **** | <0.0001 |
| HUM | 125 | *spn-A$^{null}$, polq$^{null}$, polq$^{\Delta L}$* **vs.** *spn-A$^{null}$, polq$^{null}$, POLQ$^{+}$* | ** | 0.0019 |
| HUM | 125 | *spn-A$^{null}$, polq$^{null}$, polq$^{\Delta L}$* **vs.** *spn-A$^{null}$, polq$^{null}$, polq$^{HUM}$* | *** | 0.0001 |
| HUM | 125 | *spn-A$^{null}$, polq$^{null}$, POLQ$^{+}$* **vs.** *spn-A$^{null}$, polq$^{null}$, polq$^{HUM}$* | **** | <0.0001 |
| HUM | 250 | *spn-A$^{null}$, polq$^{null}$, polq$^{\Delta L}$* **vs.** *spn-A$^{null}$, polq$^{null}$, POLQ$^{+}$* | **** | <0.0001 |
| HUM | 250 | *spn-A$^{null}$, polq$^{null}$, polq$^{\Delta L}$* **vs.** *spn-A$^{null}$, polq$^{null}$, polq$^{HUM}$* | ns | 0.9474 |
| HUM | 250 | *spn-A$^{null}$, polq$^{null}$, POLQ$^{+}$* **vs.** *spn-A$^{null}$, polq$^{null}$, polq$^{HUM}$* | **** | <0.0001 |
| HUM | 500 | *spn-A$^{null}$, polq$^{null}$, polq$^{\Delta L}$* **vs.** *spn-A$^{null}$, polq$^{null}$, POLQ$^{+}$* | ns | 0.9486 |

| | | | | |
|---|---|---|---|---|
| HUM | 500 | *spn-A*$^{null}$, *polq*$^{null}$, *polq*$^{\Delta L}$ **vs.** *spn-A*$^{null}$, *polq*$^{null}$, *polq*$^{HUM}$ | ns | >0.9999 |
| HUM | 500 | *spn-A*$^{null}$, *polq*$^{null}$, *POLQ*$^{+}$ **vs.** *spn-A*$^{null}$, *polq*$^{null}$, *polq*$^{HUM}$ | ns | 0.9486 |
| FUS | 50 | *spn-A*$^{null}$, *polq*$^{null}$, *polq*$^{\Delta L}$ **vs.** *spn-A*$^{null}$, *polq*$^{null}$, *POLQ*$^{+}$ | *** | 0.0007 |
| FUS | 50 | *spn-A*$^{null}$, *polq*$^{null}$, *polq*$^{\Delta L}$ **vs.** *spn-A*$^{null}$, *polq*$^{null}$, *polq*$^{FUS}$ | ns | 0.0974 |
| FUS | 50 | *spn-A*$^{null}$, *polq*$^{null}$, *POLQ*$^{+}$ **vs.** *spn-A*$^{null}$, *polq*$^{null}$, *polq*$^{FUS}$ | **** | <0.0001 |
| FUS | 125 | *spn-A*$^{null}$, *polq*$^{null}$, *polq*$^{\Delta L}$ **vs.** *spn-A*$^{null}$, *polq*$^{null}$, *POLQ*$^{+}$ | ** | 0.0054 |
| FUS | 125 | *spn-A*$^{null}$, *polq*$^{null}$, *polq*$^{\Delta L}$ **vs.** *spn-A*$^{null}$, *polq*$^{null}$, *polq*$^{FUS}$ | ** | 0.0012 |
| FUS | 125 | *spn-A*$^{null}$, *polq*$^{null}$, *POLQ*$^{+}$ **vs.** *spn-A*$^{null}$, *polq*$^{null}$, *polq*$^{FUS}$ | **** | <0.0001 |
| FUS | 250 | *spn-A*$^{null}$, *polq*$^{null}$, *polq*$^{\Delta L}$ **vs.** *spn-A*$^{null}$, *polq*$^{null}$, *POLQ*$^{+}$ | *** | 0.0003 |
| FUS | 250 | *spn-A*$^{null}$, *polq*$^{null}$, *polq*$^{\Delta L}$ **vs.** *spn-A*$^{null}$, *polq*$^{null}$, *polq*$^{FUS}$ | ns | 0.6181 |
| FUS | 250 | *spn-A*$^{null}$, *polq*$^{null}$, *POLQ*$^{+}$ **vs.** *spn-A*$^{null}$, *polq*$^{null}$, *polq*$^{FUS}$ | **** | <0.0001 |
| FUS | 500 | *spn-A*$^{null}$, *polq*$^{null}$, *polq*$^{\Delta L}$ **vs.** *spn-A*$^{null}$, *polq*$^{null}$, *POLQ*$^{+}$ | ns | 0.9617 |
| FUS | 500 | *spn-A*$^{null}$, *polq*$^{null}$, *polq*$^{\Delta L}$ **vs.** *spn-A*$^{null}$, *polq*$^{null}$, *polq*$^{FUS}$ | ns | >0.9999 |
| FUS | 500 | *spn-A*$^{null}$, *polq*$^{null}$, *POLQ*$^{+}$ **vs.** *spn-A*$^{null}$, *polq*$^{null}$, *polq*$^{FUS}$ | ns | 0.9617 |

Two-way ANOVA and Sidak's multiple comparisons test, *p=0.01 to 0.05, **p=0.001 to 0.01, ***p=0.0001 to 0.001, ****p=<0.0001, ns=not significant.
